# Zebrafish *hhex* null mutant develops an intrahepatic intestinal tube due to de-repression of *cdx1b* and *pdx1*

**DOI:** 10.1101/343517

**Authors:** Ce Gao, Weidong Huang, Yuqi Gao, Ji Jan Lo, Lingfei Luo, Honghui Huang, Jun Chen, Jinrong Peng

**Author notes:** These authors contributes equally to this work (Ce Gao, Weidong Huang).

## Abstract

The hepatopancreatic duct (HPD) system links the liver and pancreas to the intestinal tube and is composed of the extrahepatic biliary duct, gallbladder and pancreatic duct. Haematopoietically-expressed-homeobox (Hhex) protein plays an essential role in the establishment of HPD, however, the molecular mechanism remains elusive. Here we show that zebrafish *hhex*-null mutants fail to develop the HPD system characterized by lacking the biliary marker Annexin A4 and the HPD marker *sox9b*. The mutant HPD system is replaced by an intrahepatic intestinal tube characterized by expressing the intestinal marker *fatty-acid-binding–protein 2a* (*fabp2a*). Cell lineage analysis showed that this intrahepatic intestinal tube is not originated from hepatocytes or cholangiocytes. Further analysis revealed that *cdx1b* and *pdx1* were expressed ectopically in the intrahepatic intestinal tube and knockdown of *cdx1b* and *pdx1* restored the expression of *sox9b* in the mutant. Chromatin-immunoprecipitation analysis shows that Hhex binds to the promoters of *pdx1* and *cdx1b* genes to repress their expression. We therefore propose that Hhex, Cdx1b and Pdx1 form a genetic network governing the patterning and morphogenesis of the HPD and digestive tract systems in zebrafish.

## INTRODUCTION

Organogenesis of the zebrafish digestive system is a dynamic process that starts from 26 hours post fertilization (hpf) and finishes at around 54 hpf followed by organ growth and maturation stages (Field et al. 2003b; Zaret 2008; Goessling and Stainier 2016). During the 26-54 hpf time window, hepatoblasts and pancreatic precursor cells are first differentiated from the endodermal rod and finally form a discrete liver and pancreatic bud, respectively (Field et al. 2003b; Huang et al. 2008; Tao and Peng 2009). The liver bud, gall bladder and pancreatic bud are linked by a ductal network named as the HPD system that finally opens through the hepatopancreatic ampulla to the intestinal tract to become part of the digestive system (Dong et al. 2007; Delous et al. 2012; Manfroid et al. 2012). Extensive work has been carried out to study the organogenesis of the liver and pancreas and a number of regulatory factors have been identified (Zaret 2008; Shih et al. 2013; Goessling and Stainier 2016). In contrast, only a few factors have been identified to specify the HPD. For example, *sox9b* is found to be an essential gene for specifying HPD (Delous et al. 2012; Manfroid et al. 2012), and Fgf10 to be a key signaling molecule for patterning and differentiation of the HPD system (Dong et al. 2007). Molecular marker staining has revealed that the identity of HPD cells is clearly distinct from but is related to the hepatoblasts and pancreatic precursor cells. For example, HPD cells are characterized to be Annexin A4- and Sox9b-positive cells (Dong et al. 2007; Delous et al. 2012; Manfroid et al. 2012; Zhang et al. 2014). Although the relationship between HPD cells and hepatic or pancreatic cells in zebrafish has been investigated in several cases the relationship between HPD cells and intestinal epithelium are hardly investigated.

Hhex, also known as PRH (proline-rich homeodomain) is a homeobox-type transcriptional factor which was first identified in the avian and human haematopoietic and liver cells (Crompton et al. 1992; Hromas et al. 1993). Later studies showed that Hhex plays important roles in cell proliferation and differentiation (Soufi and Jayaraman 2008). Hhex is involved in regulating the development of blood cells (Paz et al. 2010; Goodings et al. 2015), liver (Bort et al. 2006; Rankin et al. 2011; Watanabe et al. 2014), pancreas (Bort et al. 2004; Zhao et al. 2012) and heart (Hallaq et al. 2004; Liu et al. 2014) in different species. Regarding the role of Hhex in the development of the HPD system, Bort et al found that Hhex-positive progenitors in the Hhex^LacZ/LacZ^ null mice conferred a duodenal-like cell fate (Bort et al. 2006). Hunter and colleagues observed that deletion of Hhex in the hepatic diverticulum (*Foxa3-Cre;Hhex^d2,3/−^*) resulted in a small and cystic liver together with absence of the gall bladder and the extrahepatic bile duct (Hunter et al. 2007), demonstrating that Hhex is required the hepatobiliary development in mice. Importantly, their anatomic evidence showed that the extrahepatic biliary duct is replaced by duodenum in *Foxa3-Cre;Hhex^d2,3/−^* mouse embryos (Hunter et al. 2007). Recent studies have shown that Hhex can direct the differentiation of stem cells to hepatocytes (Kubo et al. 2010; Arterbery and Bogue 2016). Hhex fulfils its function by regulating the expression of a number of downstream genes(Cong et al. 2006; Williams et al. 2008; Watanabe et al. 2014) and itself is regulated by factors including HNF3β, Smad, Wnt and Sox17 signaling etc (Denson et al. 2000; Zhang et al. 2002; Rankin et al. 2011; Liu et al. 2014). Mutation in *hhex* is associated with certain human diseases (Shields et al. 2016).

In zebrafish, morpholino-mediated gene knockdown showed that *hhex* is essential for liver and pancreas development (Wallace et al. 2001). In addition to its expression in the liver and pancreatic bud, *hhex* is also highly expressed in the HBPD precursor cells, displaying a similar dynamic expression patterns as in mice (Bogue et al. 2000; Bort et al. 2004), suggesting a role of Hhex in the development of HPD in zebrafish. In this report, by studying zebrafish *hhex* null mutants, we find that loss-of-function of Hhex leads to the formation of an intrahepatic intestinal tube at the expense of organogenesis of the HPD system. Molecular study reveals that Hhex represses the expression of *cdx1b* and *pdx1* to safeguard the cell identity and morphogenesis of the HPD system.

## Results

### Loss-of-function of *hhex* leads to abnormal liver and exocrine pancreas development

In zebrafish, hepatic and pancreatic differentiations are characterized by thickening of the foregut endoderm at around 28hpf (Ober et al. 2003; Huang et al. 2008; Tao and Peng 2009; Goessling and Stainier 2016). We performed a WISH to check the expression of *hhex* at 28hpf and found that the *hhex* transcripts were detected in the prospective liver bud and pancreatic bud (Fig. 1A). The zebrafish genome contains a single copy of the *hhex* gene located on chromosome 12 (Fig. 1B, upper panel). To generate *hhex*-knockout mutants, we designed a guide-RNA (gRNA) based on the sequence of the exon 1 of the *hhex* gene (Fig. 1B, lower panels) and co-injected *hhex*-specific gRNA with *Cas9* mRNA into one-cell stage embryos. The efficiency of *hhex*-gRNA in the injected embryos was about ∼40% (Fig. S1A). F1 progenies were screened for *hhex* mutants and two independent deletion alleles were obtained, with one deleted 17 bp (*hhex^zju1^* allele) and another 11 bp (*hhex^zju2^* allele) in the exon 1 of *hhex* (Figure 1*C*). Each of these two alleles causes a frame-shift to the open reading frame (ORF) of *hhex* and introduces an early stop codon to disrupt the translation of *hhex* (Fig. S1B), suggesting that both are likely null alleles.

**Fig. 1.**
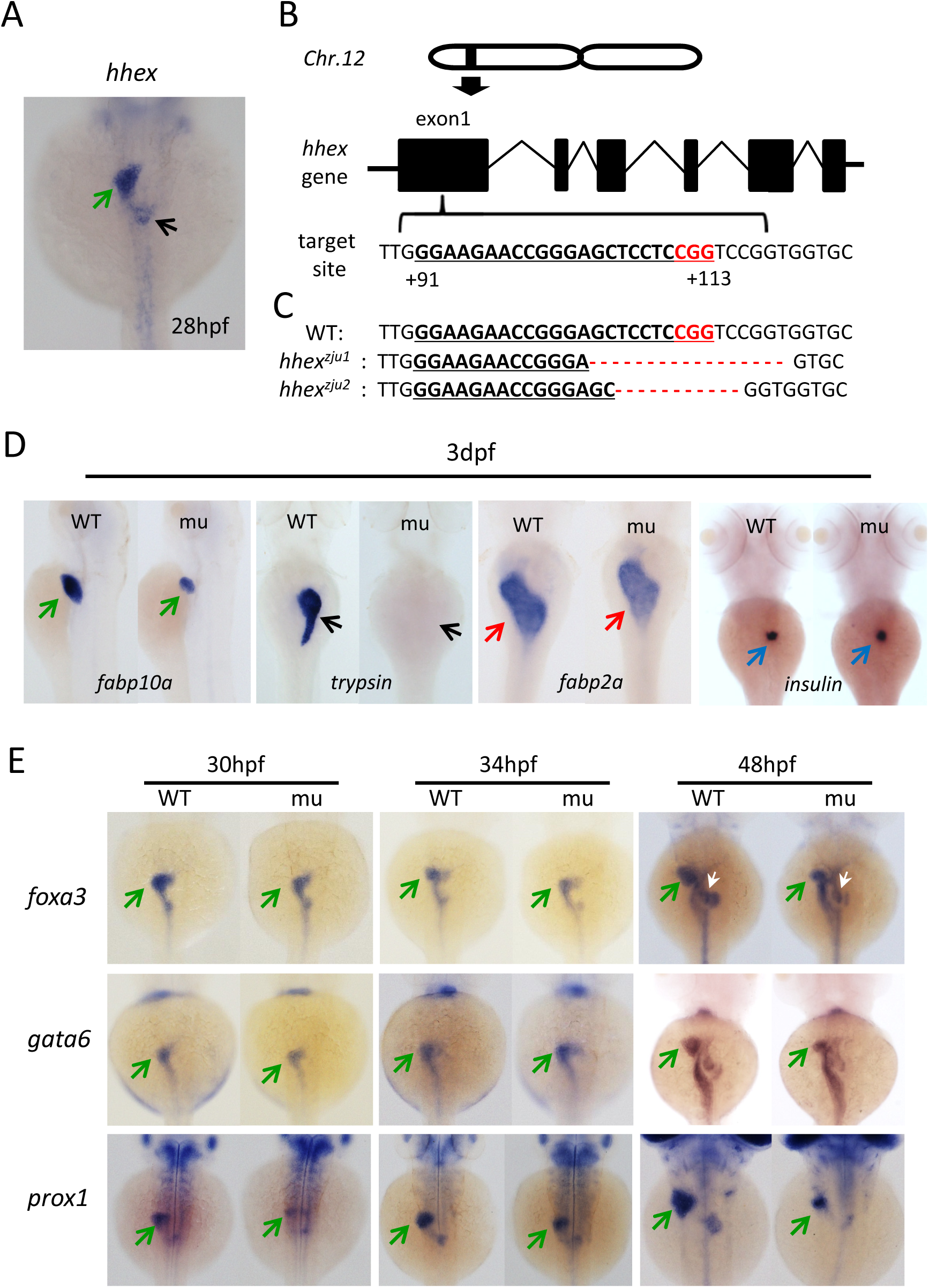
*hhex^zju1^* confers a small liver and pancreas phenotype. (A) WISH showing the expression pattern of *hhex* in a representative embryo at 28hpf. (B) Diagrams showing the chromosome position (upper panel) and genomic structure (middle panel) of *hhex*. Bottom panel: the gRNA target sequence (+91 to +113 from the start codon of *hhex*). (C) Sequence alignment showing the 17bp and 11bp deletion in *hhex^zju1^* and *hhex^zju2^*, respectively. (D) WISH using *fabp10a*, *trypsin*, *fabp2a* and *insulin* probes on embryos at 3dpf. (E) WISH using *foxa3*, *gata6* and *prox1* probes on WT and *hhex^zju1^* mutant at 30hpf, 34hpf and 48hpf. WT, wild type; mu, *hhex^zju1^*; green arrow, liver bud; black arrow, exocrine pancreas; red arrow, intestinal tube; blue arrow, islet; white arrow, swimming bladder.

To check the digestive organs development in *hhex^zju1^* we performed WISH using the *fabp10a* (liver marker), *trypsin* (exocrine pancreas marker), *fabp2a* (intestine marker) and *insulin* (islet marker) probes, on the WT and *hhex^zju1^* embryos at 3 days post-fertilization (dpf). The result showed that the *hhex^zju1^* mutant displays a small liver phenotype and is absent of a detectable exocrine pancreas (Fig. 1D) whilst the development of the intestine and islet was not obviously affected (Fig. 1D). To determine the time point when the mutant phenotype is discernable we performed WISH using pan-endodermal markers *foxa3* and *gata6* and early hepatic marker *prox1*. We found that the initiation of the liver primordium appeared to be normal in the *hhex^zju1^* mutant at 30- and 34-hpf as revealed by these molecular markers (Fig. 1E). However, by 48hpf, the growth of the liver bud appeared to be retarded in the *hhex^zju1^* mutant (Fig. 1E). The *hhex^zju2^* mutant also displayed a small liver and undetectable exocrine pancreas phenotype (Fig. S1C). Therefore, as observed in the *hhex-*conditional-knockout mice (Bort et al. 2006; Hunter et al. 2007), Hhex is essential for the outgrowth of the liver bud but not specification of the hepatic cells.

### Generation of the *Tg(bhmt:EGFP)* reporter fish that faithfully recapitulates the expression pattern of the endogenous *bhmt*

The transgenic reporter fish *Tg(fabp10a:DsRed; elastase: GFP)* is frequently used in the liver development study (Dong et al. 2007). However, due to the fact that the expression of *fabp10a* is only weakly detectable at around 48hpf (Her et al. 2003), it is desirable, especially in the case of studying liver development in *hhex^zju1^*, to obtain a reporter line that can robustly indicate the liver development before 48hpf. The *betaine homocysteine S-methyltransferase* (*bhmt*) is strongly expressed in the liver primordium in 34hpf-old zebrafish embryos (Yang et al. 2011). The zebrafish genome contains a single copy of the *bhmt* gene located on chromosome 21 (Fig. 2A, upper panel). A genomic DNA fragment 5.2kb upstream of the translation start codon ATG of *bhmt* was amplified by PCR using primer pair *bhmt* promoter (Table S1). This DNA fragment was cloned into pEGFP-1 upstream of the *EGFP* gene to generate the *bhmt:EGFP* plasmid which was then injected into one-cell stage embryos (Fig. 2A, lower panel). The *Tg(bhmt:EGFP)* transgenic fish was obtained from the progenies of the injected embryos. In *Tg(bhmt:EGFP)*, EGFP signal was detected in the liver primordium at 32hpf (Fig. 2B, outlined by dashed lines), and in the liver bud at 2-, 3-, 4-, 5- and 6-dpf. EGFP is also detected in the yolk syncytial layer (YSL), but is specifically restricted in the liver at 6dpf when the yolk is fully absorbed (Fig. 2B). In the *Tg(bhmt:EGFP)* background, we performed immunostaining of Bhmt in the cryosections obtained from the embryos at 32hpf and 2-, 4- and 6-dpf. The result showed that the EGFP signal is fully co-localized with the Bhmt signal in the liver and YSL (Fig. 2C). Therefore, the expression pattern of EGFP in *Tg(bhmt:EGFP)* faithfully recapitulates the expression pattern of the endogenous *bhmt* gene.

**Fig. 2.**
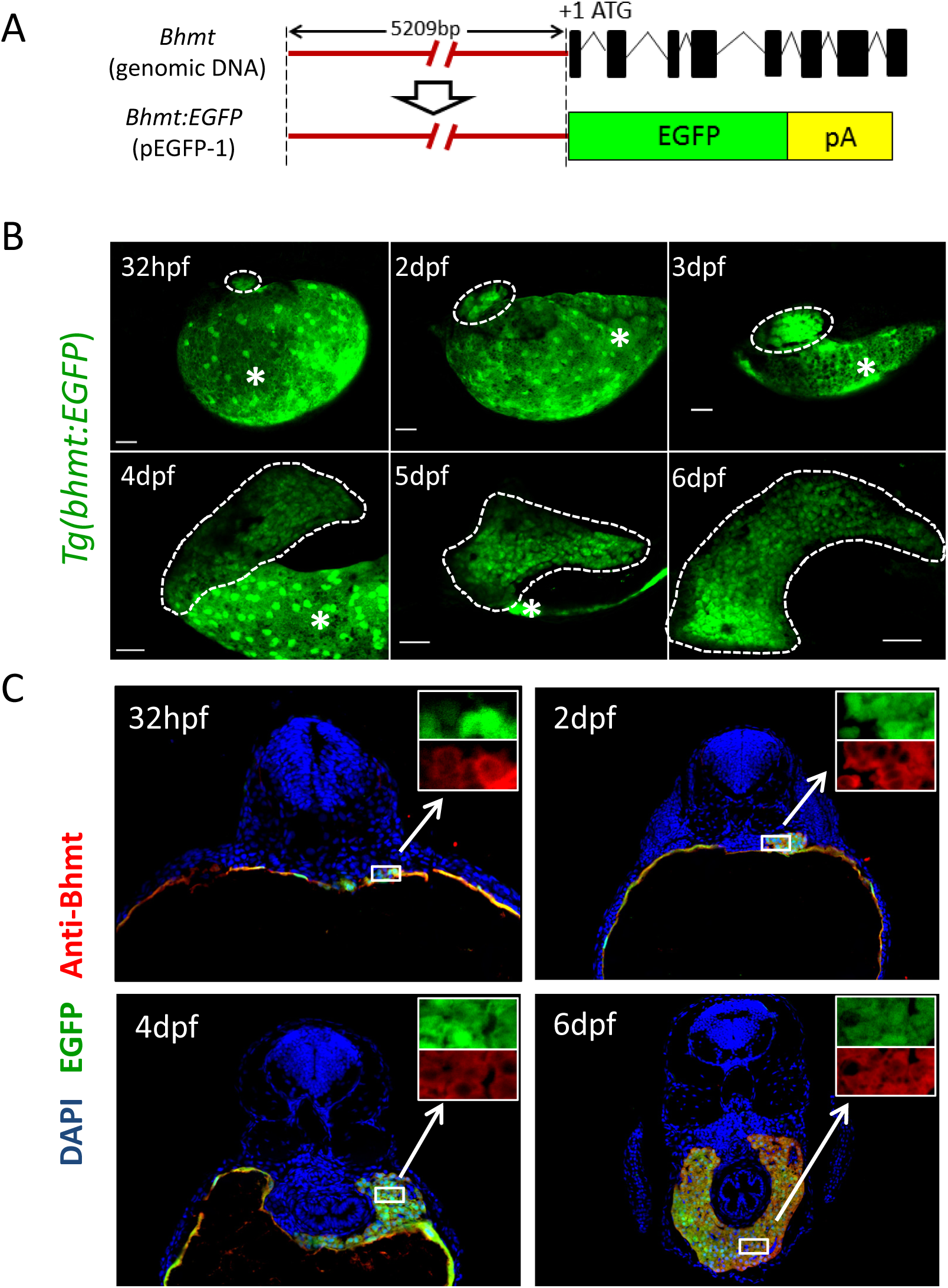
*Tg(bhmt:EGFP)* recapitulates the expression pattern of the endogenous *bhmt*. (A) Diagrams showing the genomic structure of *bhmt* (upper panel) and the *bhmt:EGFP* construct (lower panel). A 5.209kb DNA fragment upstream of the start codon of *bhmt* is cloned upstream of the reporter gene *EGFP*. (B) Examining the expression pattern of Egfp in *Tg(bhmt:EGFP)* at 32hpf, 2dpf to 6dpf. The liver region is circulated with a dashed line. *, yolk. (C) Co-immunostaining of EGFP (in green) and the endogenous Bhmt (in red) in *Tg(bhmt:EGFP)* at 32hpf, 2dpf, 4dpf and 6dpf. The insets represent the high magnification view of the boxed regions.

### *hhex^zju1^* develops an intrahepatic intestinal tube

To further characterize the small liver phenotype in *hhex^zju1^* we crossed the *hhex^zju1^* heterozygous fish (*hhex^zju1/+^*) with the *Tg(bhmt:EGFP)* reporter fish to get the *Tg(bhmt:EGFP) hhex^zju1/+^* fish. Through analyzing the cryosections from the *Tg(bhmt:EGFP) hhex^zju1^* embryos at 5dpf, we surprisingly found a lumenized structure in the mutant liver which was not present in WT (Fig. 3A). By analyzing serial sections of a 5dpf-old mutant embryo we found that this lumen structure is connected to and also opens to the main intestinal tube (Fig. 3B, highlighted by dashed lines), which was never observed in a WT embryo (Fig. S2A). Besides, the cells surrounding the lumen of *hhex^zju1^* are EGFP^−^ cells (Fig. 3A,B), excluding the hepatic nature of the cells surrounding the lumen. The presence of this lumen structure in *hhex^zju1^* is further confirmed by reconstruction of confocal images into a three-dimension (3-D) image (Fig. 3C). The *hhex^zju2^* mutant also developed an intrahepatic lumen (Fig. S2B).

**Fig. 3.**
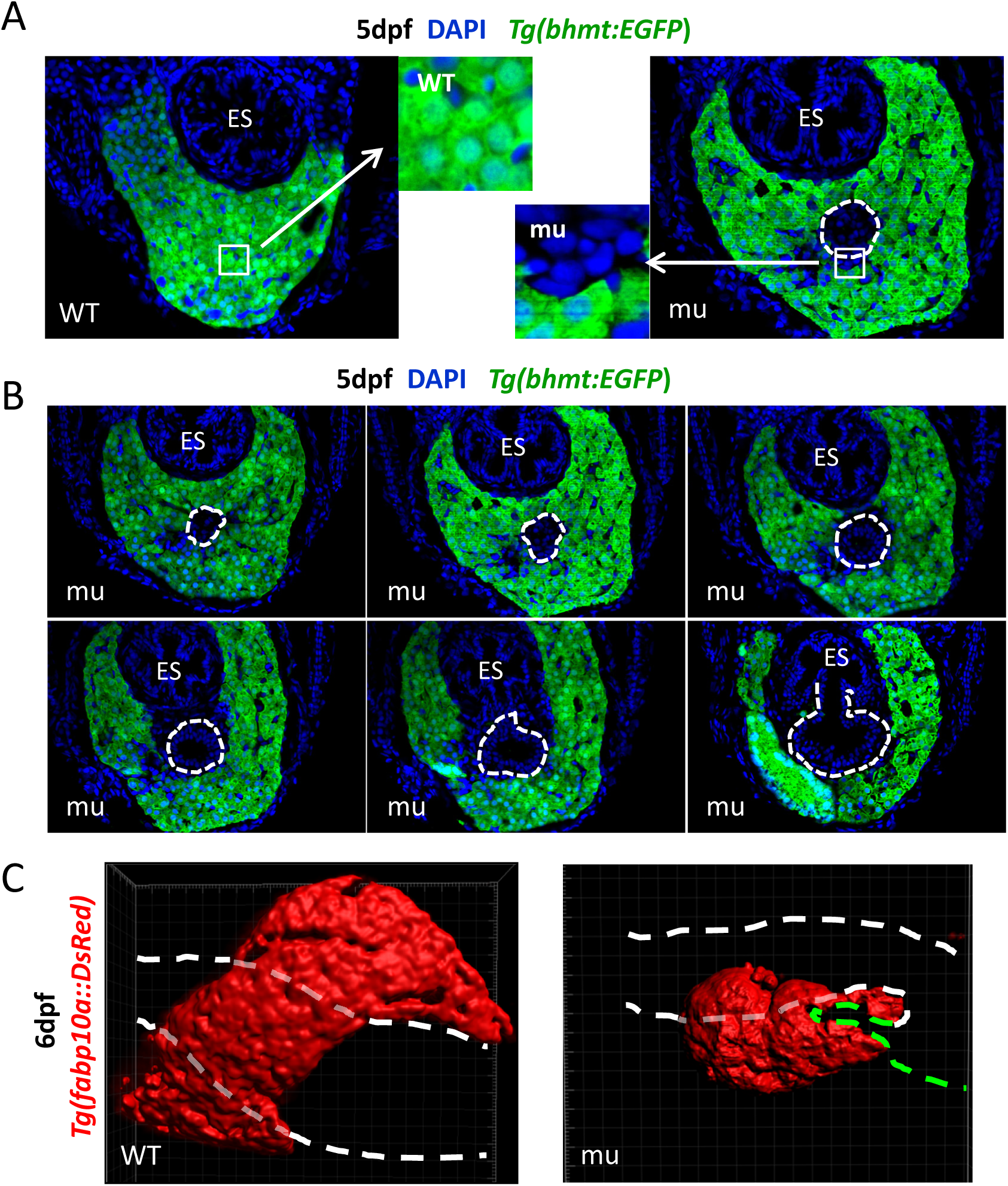
*hhex^zju1^* develops an intrahepatic lumen structure. (A) A confocal section showing the intrahepatic lumen (circulated with a white dashed line) in the *hhex^zju1^* mutant (mu) in the *Tg(bhmt:EGFP)* background. The insets represent the high magnification view of the boxed regions. (B) Consecutive confocal sections of the liver in an *hhex^zju1^* embryo (mu) in the *Tg(bhmt:EGFP)* background showing the intrahepatic lumen (highlighted with dashed lines) is finally fused to the junction region between the esophagus (ES) and intestine. (C) 3-D reconstruction of the liver in a WT and an *hhex^zju1^* embryo at 6dpf, respectively, in the *Tg(fabp10a:DsRed)* background. Pair of white dashed lines, indicating the digestive tract (esophagus plus intestine); green dashed line, indicating the intrahepatic lumen structure in *hhex^zju1^*.

We went further to determine the identity of the EGFP^−^ cells surrounding the lumen. Monoclonal antibody 2F11 specifically recognizes Annexin A4 expressed the ductal cells which can be used to mark the intrahepatic biliary ducts and the HPD system (Zhang et al. 2014). Immunostaining using 2F11 showed that the ductal system in the liver (Fig. 4A,B: 2dpf-L, 3dpf-L, 5dpf-L and 7dpf-L) and the gall bladder (Fig. 4A,B: 5dpf-R and 7-dpf-R) in WT at 3-, 5- and 7-dpf is clearly visible. However, in the *hhex^zju1^* mutant, the intrahepatic biliary ducts are severely compromised (Fig. 4A,B: 2dpf-L, 3dpf-L, 5dpf-L and 7dpf-L) and the HPD system (including the gallbladder) are not formed (Fig. 4A,B: 5dpf-R and 7-dpf-R). This result demonstrates that the lumen structure is not an outcome of over-expansion of the ductal system.

**Fig. 4.**
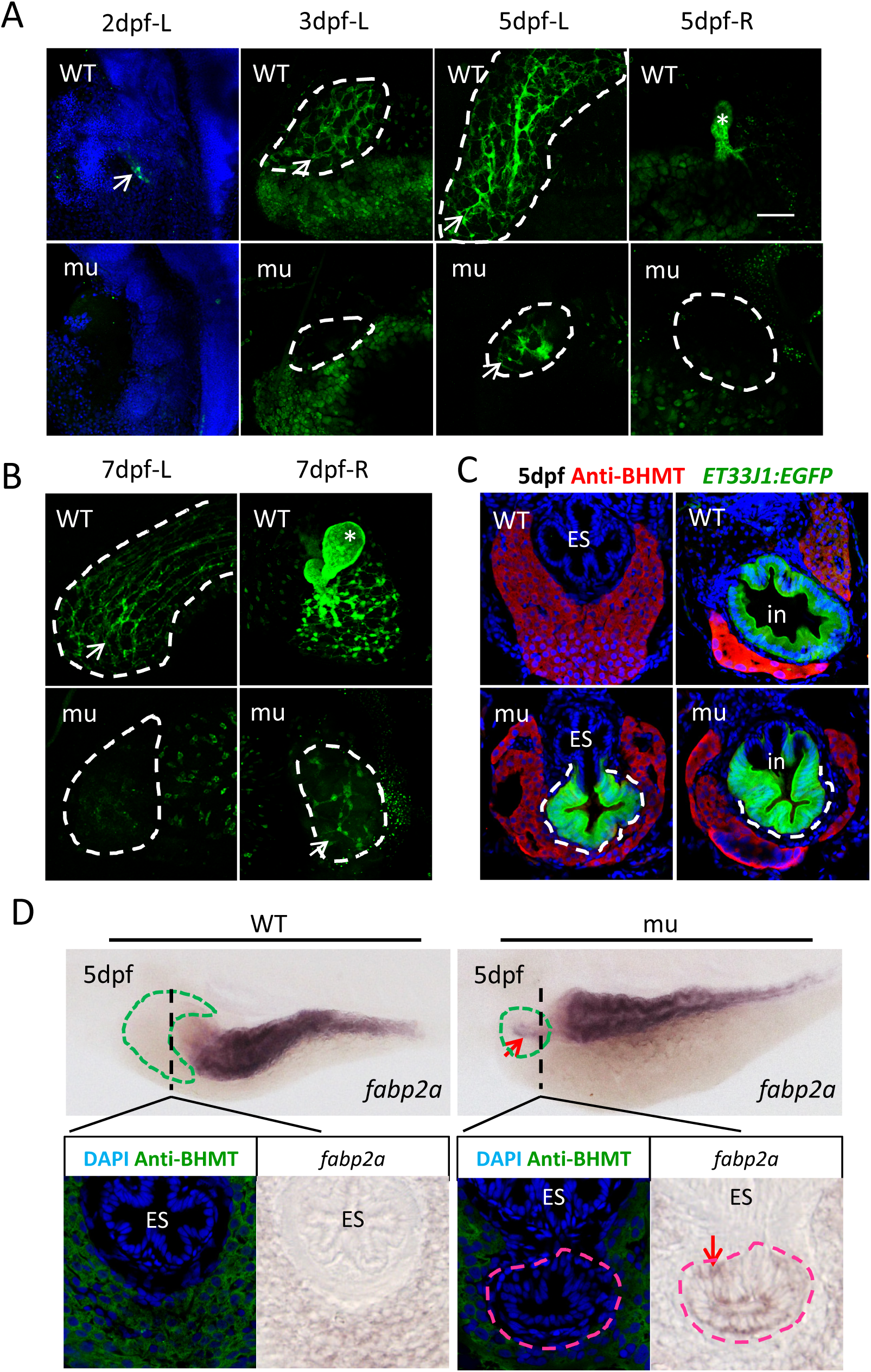
*hhex^zju1^* develops an intrahepatic intestinal tube. (A,B) Immunostaining using the 2F11 antibody to compare the HBPD development between the WT and *hhex^zju1^* mutant at 2dpf, 3dpf, 5dpf and 7dpf (dpf-L: left side of the embryo, dpf-R: right side). The liver region is circulated with a white dashed line. white arrow, bile duct system; *, gallbladder. (C) Confocal sections of the liver in a WT and an *hhex^zju1^* embryo (mu) in the *Tg(ET33J1:EGFP)* background showing the cells surrounding the intrahepatic lumen (highlighted with dashed lines) were EGFP-positive at 5dpf. Note that the posterior part of the esophagus (ES) was EGFP-negative in both WT and mutant. (D) Upper panels: WISH using the intestinal marker *fabp2a* on the WT and *hhex^zju1^* mutant (mu) embryos at 5dpf. The liver is circulated with a green dashed line. A red arrow points to the intrahepatic lumen which is *fabp2a*-positive. Lower panels: After WISH using the *fabp2a* probe, embryos were sectioned through the plane as indicated by the black dashed line and stained with the Bhmt antibody. The intrahepatic lumen in *hhex^zju1^* is of the nature of *fabp2a*-positive cells.

The zebrafish enhancer trapping line *Tg(ET33J1:EGFP)* (http://plover.imcb.a-star.edu.sg/webpages/ET33-J1.html) strongly expresses the reporter EGFP in the gut epithelia together with weak expression in the anterior part of the esophagus (Fig. S3). We crossed *hhex^zju1/+^* with the *Tg(ET33J1:EGFP)* and analyzed the EGFP signal in the *hhex^zju1^* background. As expected, in the WT embryo at 5dpf, the EGFP signal was strongly expressed in the intestinal bulb, but was not observed in the liver and the posterior part of the esophagus connecting to the intestinal bulb (Fig. 4C, upper panels). Interestingly, in *hhex^zju1^*, the intrahepatic lumen displayed a strong EGFP signal that finally fused with the digestive tract at the junction between the esophagus (which is EGFP^−^) and the intestinal bulb (Fig. 4C, lower panels).

The expression of *fabp2a* is normally restricted in the intestine. Although the expression pattern of *fabp2a* in *hhex^zju1^* appears to be similar to that in WT at 3dpf (Fig. 1D), careful examination of the *fabp2a* stained 5dpf-old *hhex^zju1^* embryos revealed that there is a short protrusion of *fabp2a*-positive cells (Fig. 4D, upper panels), which looks to match the position of the lumenized structure revealed by 3-D reconstruction (Fig. 3C). We then sectioned the *fabp2a*-WISH embryos and found that the cells forming the lumenized structure in the *hhex^zju1^* mutant liver were indeed of being *fabp2a*-positive (Fig. 4D, lower panels). Therefore, the lumenized structure is an intrahepatic intestinalized tube.

### The intrahepatic intestine is not originated from Fabp10a- or TP1-positive cells

In the transgenic fish *Tg(fabp10a: CreERT2)*, the expression of *CreERT2* is driven by the hepatocyte specific promoter *fabp10a*. Introducing *Tg(fabp10a: CreERT2)* to the *Tg(β-actin:loxP-DsRed-STOP-loxP-GFP)* background will, upon the tamoxifen treatment, genetically label all embryonic hepatocytes to express EGFP permanently (Gao et al. 2018). We first crossed *hhex^zju1/+^* with *Tg(β-actin: loxP-DsRed-STOP-loxP-GFP)* and *Tg(fabp10a: CreERT2)*, respectively, and then crossed *hhex^zju1/+^ Tg(fabp10a: CreERT2)* fish with *hhex^zju1/+^ Tg(β-actin:loxP-DsRed-STOP-loxP-GFP)* fish. As expected, all hepatocytes were labeled by EGFP (EGFP^+^) in either *Tg(fabp10a: CreERT2; β-actin: loxP-DsRed-STOP-loxP-GFP)* (WT background) or *hhex^zju1^ Tg(fabp10a: CreERT2; β-actin: loxP-DsRed-STOP-loxP-GFP)* (mutant background) 5dpf-old embryos after the tamoxifen treatment at 36hpf (Fig. 5A). Examining the liver in the 5dpf-old *hhex^zju1^ Tg(fabp10a: CreERT2; β-actin: loxP-DsRed-STOP-loxP-GFP)* embryos after the tamoxifen treatment at 36hpf we found that the cells forming the intrahepatic intestinal tube in *hhex^zju1^* were EGFP-negative cells (Fig. 5A, right image panels).

**Fig. 5.**
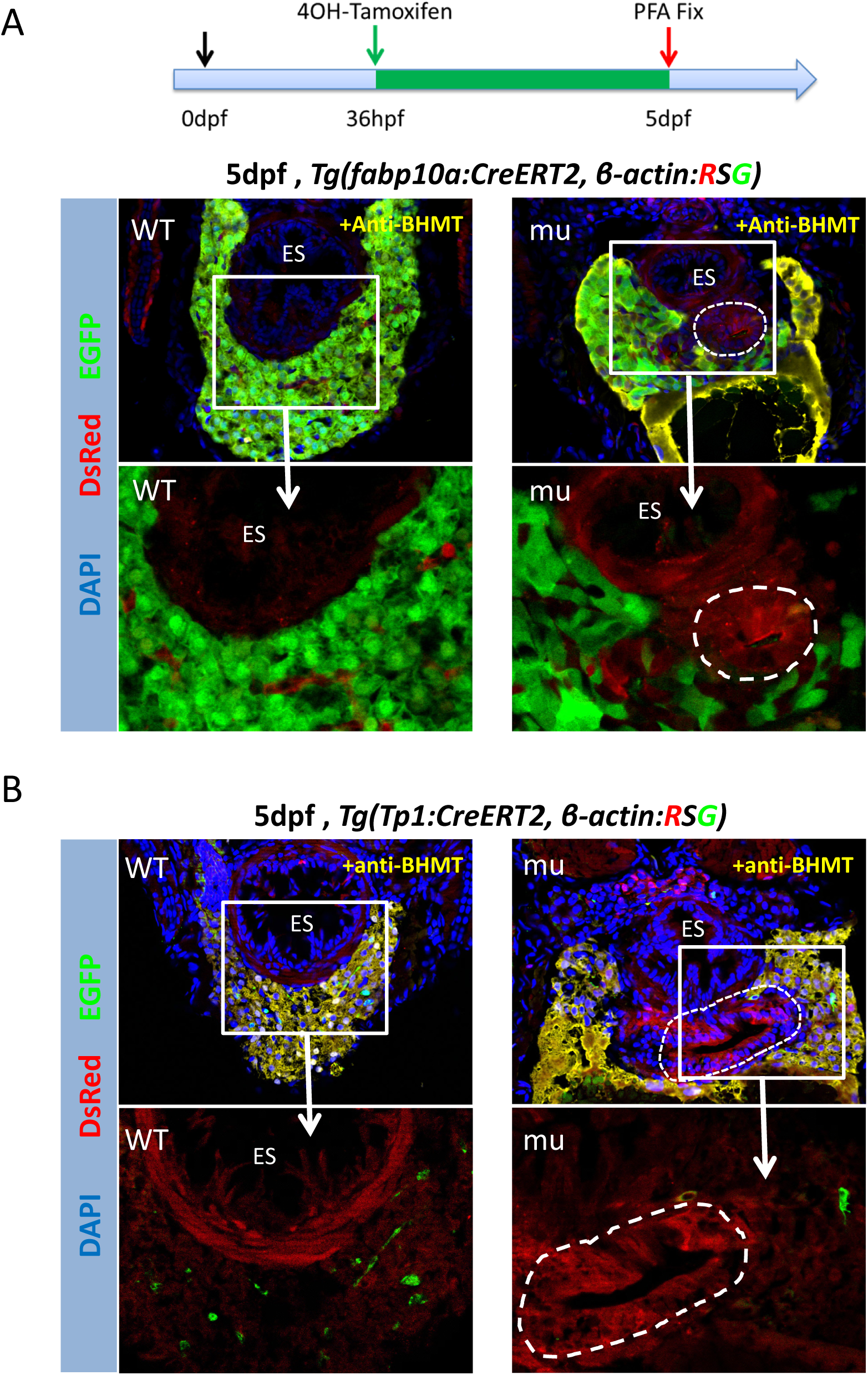
The intrahepatic intestinal tube is not originated from Fabp10a- or TP1-positive cells. (A,B) Upper panels: Co-immunostaining of Bhmt (yellow), DsRed (red) and EGFP (green) to examine the liver in the 5dpf-old WT and *hhex^zju1^* (mu) embryos in the background of *Tg(fabp10a: CreERT2;β-actin: loxP-DsRed-STOP-loxP-GFP)* (for labeling hepatocytes) (A) or of *Tg(Tp1: CreERT2; ß-actin:loxP-DsRed-STOP-loxP-GFP)* (for labeling bile duct cells) (B) after the tamoxifen treatment at 36hpf. Lower panels: The high magnification views of the boxed regions in (A,B) show that the cells forming the intrahepatic intestinal tube (circulated with a white dashed line) were EGFP-negative. DAPI was used to stain the nuclei. ES, esophagus.

In *Tg(TP1: CreERT2)*, the *Cre-ERT2* is expressed in the bile duct epithelia cells which can be adopted to trace the bile duct cell lineage(He et al. 2014) (Fig. 5B, left panels). We crossed *hhex^zju1/+^* with *Tg(β-actin: loxP-DsRed-STOP-loxP-GFP)* and *Tg(TP1: CreERT2)*, respectively. We then crossed *hhex^zju1/+^ Tg(TP1: CreERT2)* fish with *hhex^zju1/+^ Tg(β-actin: loxP-DsRed-STOP-loxP-GFP)* fish. Treating the progenies derived from *hhex^zju1/+^ Tg(Tp1: CreERT2; β-actin: loxP-DsRed-STOP-loxP-GFP)* with tamoxifen permanently labeled the bile duct cells as EGFP^+^ (Fig. 5B). We failed to identify any EGFP^+^ cells within the intrahepatic intestinal tube in *hhex^zju1^* at 5dpf after tamoxifen treatment at 36hpf (Fig. 5B, right panels). Therefore, the cells surrounding the intrahepatic intestinal tube are originated neither from the differentiated hepatocytes nor form the bile duct cells.

### The intrahepatic intestine is formed at the expense of the HPD system

We have shown that *hhex* is expressed in the prospective liver and pancreatic buds at 28hpf (Fig. 1A). To understand how Hhex is involved in the HPD development, we checked the temporal and spatial expression patterns of the *hhex* transcripts at 24 and 28hpf by WISH. The result showed that the expression of *hhex* was detected in the HPD cells at 28hpf in the WT embryos (Fig. 6A), suggesting that Hhex likely plays a role in the development of the HPD system as well. Notably, WISH using the *hhex* probe showed that the *hhex^zju1^* mutant produced *hhex* transcripts in the liver bud (Fig. 6A), suggesting that the transcription of *hhex* in the liver bud is not affected by the *hhex^zju1^* mutation. However, the *hhex* transcripts were almost abolished in the region prospected for the exocrine pancreas in the *hhex^zju1^* mutant at 28hpf (Fig. 6A), which might explains why the *hhex^zju1^* mutant failed to develop an exocrine pancreas (Fig. 1D).

**Fig.6.**
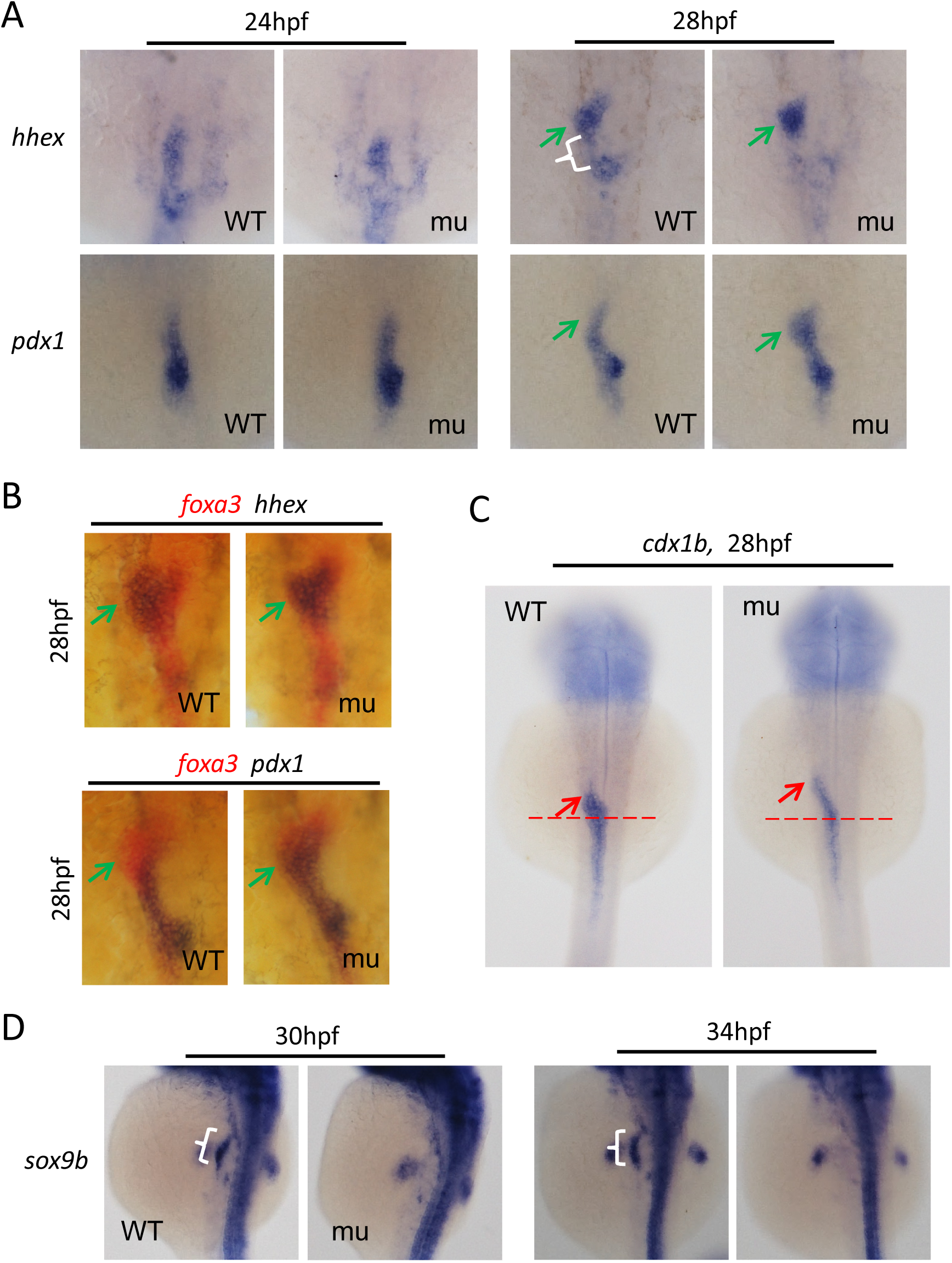
Loss-of-function of Hhex causes an ectopic expression of *cdx1b* and *pdx1* and abolishes the *sox9b* expression in the prospective HPD region at 28hpf. (A-D) WISH using the *hhex* or *pdx1* probe (A), *foxa3* and *hhex* or *foxa3* and *pdx1* double probes (B), *cdx1b* probe (C) or *sox9b* probe on WT and *hhex^zju1^* mutant (mu) embryos at different stages as shown. The *pdx1* expression domain overlapped with the *hhex*-positive domain only in the mutant but not in the WT at 28hpf (B). The *cdx1b* expansion domain is extended anteriorly in the mutant at 28 hpf (C). *sox9b* transcripts were undetectable in the HPD area but were clearly visible in other regions including the fin bud at 30 and 34hpf in the mutant (D). Bracket, prospective HPD; green arrow, liver; red arrow, the anterior edge of the intestinal bulb, red dashed line, left bending point for the endoderm rod.

We asked how the intrahepatic intestinal tube is formed in *hhex^zju1^*. *pdx1* (*pancreatic and duodenal homeobox 1*) is a transcription factor necessary for intestinal tube differentiation and pancreatic development in zebrafish(Yee et al. 2001). WISH using *pdx1* probe shows that there is an ectopic expression of *pdx1* in the prospective liver bud area at 28hpf but not 24hpf in *hhex^zju1^* (Fig. 6A). *foxa3* is a pan-endoderm marker. Comparing the signal patterns from WISH using the *foxa3* and *hhex* or *foxa3* and *pdx1* double probes revealed that the *pdx1* signal overlapped with the *hhex* signal only in mutant liver bud but not the WT (Fig. 6B). The zebrafish *caudal-related homeobox 1b* (*cdx1b*) gene is a specific marker for the foregut region and is essential for the normal development of the digestive tract in zebrafish (Cheng et al. 2008). WISH using the *cdx1b* probe showed that there is a protrusion of the *cdx1b*-positive domain at the tip of the foregut in *hhex^zju1^* at 28hpf (Fig. 6C).

The transcription factor gene *sox9b* has recently been identified to be a specific marker and also an essential gene for the development of the HPD system (Delous et al. 2012; Manfroid et al. 2012). We performed a WISH using the *sox9b* probe. As expected, *sox9b* is expressed in the HPD cells at 30hpf, 34hpf, 38hpf and 40hpf (Fig. 6D; Fig. S4A). However, the *sox9b* transcripts were undetectable in the prospected HPD cells in the *hhex^zju1^* mutant (Fig. 6D; Fig. S4A), demonstrating that the *hhex^zju1^* mutant fails to develop the HPD system.

To further confirm the expression patterns of *sox9b*, *cdx1b* and *pdx1* in *hhex^zju1^* embryos, we performed WISH at later embryonic stage. The results showed that in the *hhex^zju1^* mutant the *sox9b* transcripts were still undetectable at 2dpf and 3dpf in the prospected HPD cells (Fig. 7A). With the reference to the somite, WISH using the *cdx1b* probe showed that there is an obvious protrusion of the *cdx1b*-positive domain at the tip of the foregut in *hhex^zju1^* at 2-, 3-, 4- and 5-dpf (Fig. 7A; Fig. S4B). At 2 and 3dpf, *pdx1* is expressed in the domains of the posterior part of the esophagus, anterior part of the intestinal tube and pancreatic duct/bud in WT (Fig. 7A). In contrast, in *hhex^zju1^*, while the *pdx1*-positive domains in the intestinal tube and islet appeared not to be affected the domain for pancreatic duct is missing at 2dpf (Fig. 7A), which nicely explains the pancreatic phenotype in *hhex^zju1^*. Strikingly, a new *pdx1*-positive domain is formed on the left side of the intestinal tube in *hhex^zju1^* and this domain apparently extended into the liver region (Fig. 7A).

**FIG. 7.**
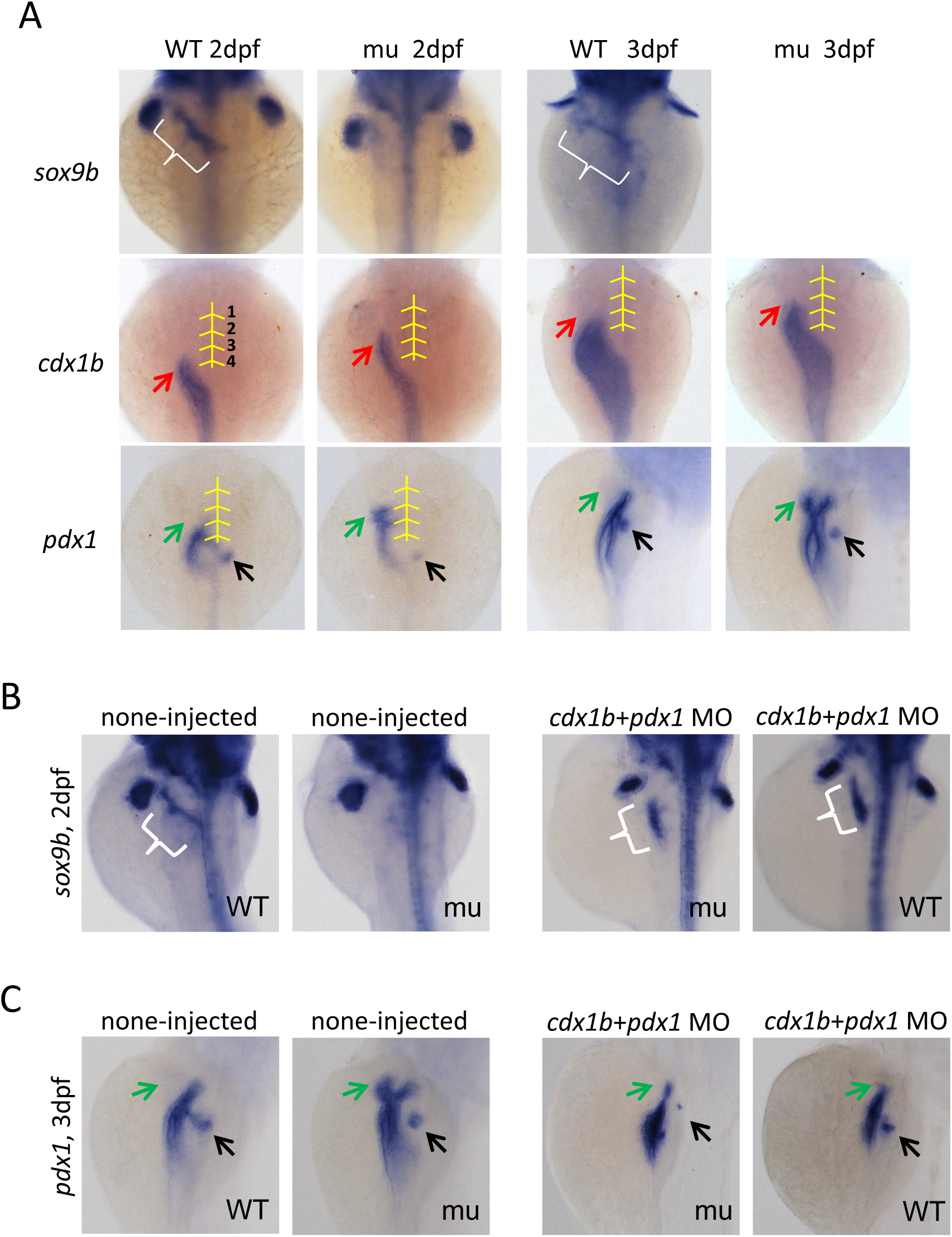
Co-injection of *cdx1b* and *pdx1* specific morpholinos restores the *sox9b* expression and abolishes the ectopic *pdx1* expression in the mutant HPD region. (A) WISH using the *sox9b*, *cdx1b* and *pdx1* probes on WT and *hhex^zju1^* mutant (mu) embryos at 2dpf and 3dpf. Abolishment of *sox9b* expression and ectopic expression of *cdx1b* and *pdx1* (the middle line and somite 1 to 4 were indicated with yellow lines.) became more evident in the mutant at 2 and 3dpf. (B) Co-injection of *cdx1b* and *pdx1* specific morpholinos restored the *sox9b* expression in the mutant HPD at 2dpf. (C) Co-injection of *cdx1b* and *pdx1* specific morpholinos resulted in disappearance of *pdx1* ectopic expression in the mutant HPD. Bracket, prospective HPD; green arrow, liver; red arrow, the anterior edge of the intestinal bulb.

We then performed WISH using *cdx1b* and *pdx1* double probes on embryos at 2-, 3- and 4-dpf. The result showed that the *cdx1b* signal nicely overlapped with the *pdx1* signal along the extra domain of the digestive tract in *hhex^zju1^* which was never observed in a WT (Fig. S4C). Furthermore, WISH using *hhex* and *pdx1* or *hhex* and *cdx1b* double probes revealed that the *pdx1* or *cdx1b* signal never overlapped with the *hhex* signal along the intestinal tube (*pdx1* overlapped with *hhex* in the islet) in a WT embryo (Fig. S4D). In contrast, in the *hhex^zju1^* mutant, the *hhex* signal (*hhex^zju1^* transcripts) overlapped with *pdx1* and *cdx1b* in the extra domain of the digestive tract (Fig. S4D).

Next, we asked whether the ectopic expression of *cdx1b* and *pdx1* was at the expense of the HPD ells. To confirm this hypothesis, we co-injected *cdx1b* and *pdx1* specific morpholinos into the WT and *hhex^zju1^* embryos to knock down the expression of these two genes. We observed that the expression of *sox9b* was restored in *hhex^zju1^* at 48hpf (Fig. 7B). Furthermore, WISH using *pdx1* showed that co-injection of *cdx1b* and *pdx1* morpholinos abolished the ectopically expressed *pdx1* in *hhex^zju1^* at 3dpf (Fig. 7C). These results suggest that the loss-of-function of *hhex* leads to the ectopic expression of *pdx1* and *cdx1b* in the HPD cells that finally developed into an intrahepatic intestine tube. Therefore, we hypothesize that Hhex determines the HPD cells fate by repressing the expression of *cdx1b* and *pdx1*.

### Hhex represses the expression of *cdx1b* and *pdx1* to safeguard the development of the HPD system

The lack of the *sox9b*-positive cells in *hhex^zju1^* suggests that Hhex is essential for the differentiation of the HPD cells. Previous studies have shown that Hhex acts either as an activator or a repressor to regulate the expression of its downstream genes (Soufi and Jayaraman 2008). The ectopic expression of *cdx1b*- and *pdx1*-positive cells in *hhex^zju1^* suggests that Hhex might negatively regulate the expression of these two genes so that to prevent the differentiation of the HPD cells to gut epithelia. To determine the relationship between Hhex and *cdx1b* or *pdx1* expression we obtained a 3,481bp and a 3,764bp genomic DNA fragment upstream of the translation start site ATG of *cdx1b* and *pdx1*, and cloned them to the upstream of the *EGFP* reporter gene to get *cdx1b:EGFP* and *pdx1:EGFP*, respectively (Fig. 8A). The *cdx1b:EGFP* and *pdx1:EGFP* plasmids were used to transfect HCT116 (Human colorectal carcinoma) cells, respectively. Examination of the EGFP fluorescence (Fig. S5A,B) and western blot analysis of the EGFP protein (Fig. 8B) showed that both plasmids achieved the expression of the *EGFP* gene, suggesting that the DNA fragments cloned harbored the promoter activity for the expression of *cdx1b* and *pdx1*, respectively.

**Fig. 8.**
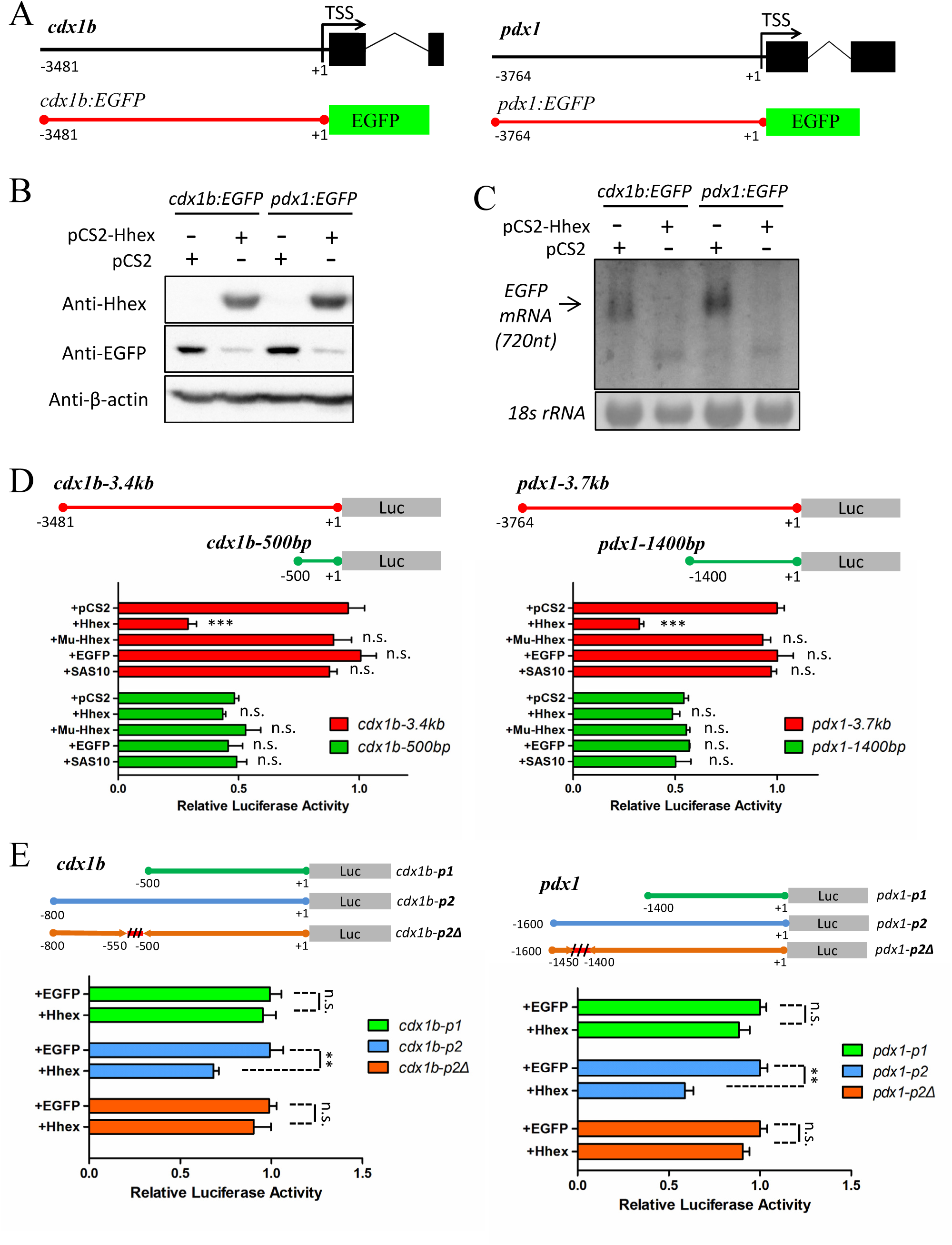
Hhex represses the transcriptional activity of *cdx1b* and *pdx1* promoters. (A). Diagram showing the promoter fragment of *cdx1b* (3.481kb, left panel) and *pdx1* (3.764kb, right panel) and their corresponding reporter construct *cdx1b:EGFP* and *pdx1:EGFP*. TSS: translation start site. (B,C). Western blotting of EGFP protein (B) and northern blotting of the *egfp* transcripts (C) in the *cdx1b:EGFP* or *pdx1:EGFP* plasmid transfected HCT116 cells with or without the pCS2-Hhex plasmid. (D). The *cdx1b* promoter regions −1 to −3481bp (*cdx1b-3.4kb*) and −1 to −500bp (*cdx1b-500bp*) (left panels) and the *pdx1* promoter regions −1 to −3764bp (*pdx1-3.7kb*) and −1 to −1400bp (*pdx1-1400bp*) (right panels) were cloned to the pGL3 vector, respectively. Luciferase activity assays showed that Hhex, but not EGFP, SAS10 or *hhex^zju1^* mutant, repressed the transcriptional activity of the *cdx1b-3.4kb* and *pdx1-3.7kb* but not of the *cdx1b-500* and *pdx1-1400bp*. (E). Three *cdx1b* promoter reporter plasmids, *cdx1b-p1*(−1 to −500bp), *cdx1b-p2* (−1 to −800bp), *cdx1b-p2Δ* (deleted −500 to −550bp region in *cdx1b-p2*) (left panels), and three *pdx1* promoter reporter plasmids, namely *pdx1-p1*(−1 to −1400bp), *pdx1-p2* (−1 to −1600bp) and *pdx1-p2Δ* (deleted −1400 to −1450 region in *pdx1-p2*) (right panels), were constructed. Luciferase activity assays show that the sequences between −500 to −550 in *cdx1b-p2* and −1400 to −1450bp in *pdx1-p2* contain the Hhex-responsive elements. All luciferase activity was normalized with Renilla luciferase vectors (phRL-TK) as the reference. * *, p<0.01; * * *. P<0.001; n.s., no significance.

We co-transfected the *pCS2-CMV:hhex* plasmid with either *cdx1b:EGFP* or *pdx1:EGFP*. Hhex was robustly expressed in the transfected HCT116 cells (Fig. 8B). Analyzing EGFP-fluorescence and -protein showed that, compared to the cells transfected with the *pCS2*-vector plus *cdx1b:EGFP* or *pCS2*-vector plus *pdx1:EGFP*, expression of Hhex suppressed the expression of EGFP (Fig. 8*B*; Fig. S5) by transcriptionally downregulating the expression of *EGFP* (Fig. 8C). To better quantify the effect of Hhex on the promoter activity of *cdx1b* and *pdx1*, we cloned the reporter gene *Firefly luciferase* (*luc*) downstream of the *cdx1b* (*cdx1-3.4kb: luc*) and *pdx1* (*pdx1-3.7kb: luc*) promoter, respectively (Fig. 8D, upper panels). Co-transfection analysis showed that Hhex, but not EGFP, Sas10 (a nucleolar protein) (Wang et al. 2016), or *hhex^zju1^* mutant, strongly suppressed the expression of Luc in *cdx1b-3.4kb: luc* and *pdx1-3.7kb: luc* (Fig. 8D, lower panels).

Hhex is reported to bind to a consensus sequence 5’-YWATTAAR-3’ (Crompton et al. 1992) By searching the promoter sequences we identified four and five putative Hhex-binding consensus sequences in the promoters for *cdx1b* and *pdx1*, respectively (Fig. S6A). However, deleting all four motifs in *cdx1b* (*cdx1b-ΔHRE1-4*) or all five motifs in *pdx1* (*pdx1-ΔHRE1-5*) did not alter the repressive effect of Hhex on the *pdx1* and *cdx1b* promoters (Fig. S6B). To determine the Hhex-responsive elements in the *pdx1* and *cdx1b* promoters, we performed ChIP using protein samples extracted from the *pCS2-HA-Hhex* plus *pdx1: luc* or *pCS2-HA-Hhex* plus *cdx1b: luc* co-transfected HCT116 cells. ChIP products were analyzed by qPCR using specific primers (Table S2) covering every 200bp of the *pdx1* 3,764bp and *cdx1b* 3,481bp DNA fragment (Fig. S6C, upper panels). Two regions in *cdx1b* promoter (−432 to −665 and –3262 to –3481) and two regions in *pdx1* promoter (−1282 to −1490 and –1950 to –2167) were identified to be enriched by the HA-tag antibody (Fig. S6C, lower panels).

We then generated a series of promoter deletion constructs and found that the −1 to −500bp region in *cdx1b* and the −1 to −1400bp region in *pdx1* have lost responses to the repressive effect of Hhex on the *luc* expression (Fig. 8D), suggesting that these regions do not harbor the Hhex-responsive elements. On the other hand, the −1 to −600bp or to −800bp of *cdx1b* and −1 to −1500bp or to −1600bp of *pdx1* exhibited strong response to the repressive effect of Hhex on the *luc* expression (Fig. S7A), which is consistent with the ChIP analysis (Fig. S6C). We therefore focused on the −1 to −600bp region in *cdx1b (cdx1b-p2)* and −1 to −1500bp region in *pdx1 (pdx1-p2)*. We generated different internal deletion constructs and found that deleting the −500 to −550bp region in *cdx1b-p2* and the −1400 to −1450bp region in *pdx1-p2* lost responsiveness to the repressive effect of Hhex on the luciferase activity (Fig. 8E). ChIP-PCR analysis showed that, compared to *cdx1b-p2* and *pdx1-p2*, Hhex is indeed no longer enriched in *cdx1b-p2Δ* or *pdx1-p2Δ* (Fig. S7B).

## DISCUSSION

In zebrafish, *hhex* is expressed in the foregut endoderm region at 26hpf (Ober et al. 2006), and then extended to the liver, pancreas and HPD cells at 48hpf (Huang et al. 2008). Importantly, zebrafish *hhex* null mutant can survive up to 7dpf. Considering the fact that the morphogenesis of the zebrafish digestive organs finishes by 52 hpf, the *hhex* null mutant provides an ideal model to study the role of Hhex not only in the liver and pancreas development but also the HPD development.

We showed here that the zebrafish *hhex* null mutant *hhex^zju1^* develops a small liver, suggesting that Hhex is not necessary for liver specification but is essential for liver bud growth as that observed in mice (Bort et al. 2004). The *hhex^zju1^* mutant fails to develop a detectable exocrine pancreas, which coincides with its role in the development of ventral pancreas in mouse (Bort et al. 2004). The *hhex^zju1^* mutant also lacks the HPD system as that reported in mice (Bort et al. 2006; Hunter et al. 2007). In contrast, loss-of-function of Hhex does not affect the overall development of the intestine. Therefore, Hhex is a key factor controlling the organogenesis of the axillary digestive organs including liver, pancreas and the HPD system but not the main digestive tract.

Anatomic analysis revealed that *hhex^zju1^* develops an intrahepatic lumen that is fused to the main intestinal tract. Cell lineage tracing study showed that this intrahepatic lumen is not originated from the hepatocytes or bile duct cells. Analysis of the expression of the intestinal marker *fabp2a* revealed that the cells forming the intrahepatic lumen were the nature of intestinal epithelia. Therefore, *hhex^zju1^* develops an intrahepatic intestinal tube. This phenotype can be explained by the fact that *cdx1b* and *pdx1*, two genes essential for the digestive tract organogenesis, are expressed not only in the normal intestinal epithelium but also ectopically in the intrahepatic lumen in *hhex^zju1^*.

Considering the fact that *sox9b* signal was not detected in the prospected HPD region in the *hhex^zju1^* mutant it is reasonable to speculate that the intrahepatic intestinal tube is originated from the HPD precursor cells. There are four lines of evidences to support this hypothesis. Firstly, it has been shown that the Hhex progenitors conferred a duodenal fate in the *Hhex*-null mice (Bort et al. 2006). In addition, the extrahepatic biliary duct is almost replaced by duodenum in the mice with conditional knockout of Hhex in the early endoderm cells (Hunter et al. 2007). Therefore, it appears that, by default, the endoderm lacking Hhex is destined to the duodenal fate. Secondly, WISH revealed that most of the *hhex*-expression domain becomes *pdx1*-positive at 28hpf in the *hhex^zju1^* mutant. Our data is consistent with the previously reported relationship between the expression patterns of *hhex* and *pdx1* (Chung et al. 2008; Xu et al. 2016). Thirdly, we show that Hhex directly repress the promoter activities of *cdx1b* and *pdx1* genes. Therefore, it is reasonable to propose that the expanded population of *pdx1*-positive cells yields the intrahepatic intestinal tube in the *hhex^zju1^* mutant. Finally, we found that knockdown of *cdx1b* and *pdx1* restored the expression pattern of *sox9b* in the prospected HPD domain in the *hhex^zju1^* mutant, further confirming the fate conversion of the HPD cells to the intestinal epithelial cells. This data also suggests that the *sox9b* gene is not a direct target of Hhex.

Based on the above, we propose a genetic network that orchestrates the organogenesis of the axillary digestive organs including liver, exocrine pancreas and the HPD system. Hhex is expressed in the liver and pancreas buds and in the HPD precursor cells. The fact that loss-of-function of Hhex affects the development of these organs suggests that Hhex functions as a positive regulator controlling the organogenesis of these organs. Sox9b is expressed in the HPD precursor cells in WT (Delous et al. 2012; Manfroid et al. 2012), however, the expression of *sox9b* is absent in the HPD precursor cells in *hhex^zju1^*, suggesting that Hhex genetically acts upstream of Sox9b although the exact relationship between Hhex and Sox9b remain to be elucidated. In a WT embryo, both Cdx1b and Pdx1 are expressed in the foregut region of the endoderm but not in the HPD precursor cells (Field et al. 2003a; Cheng et al. 2008). In the WT HPD precursor cells, Hhex suppresses the expression of *cdx1b* and *pdx1* so that to safeguard the HPD cell fate. In *hhex^zju1^*, due to the absence of Hhex, the HPD precursor cells express Cdx1b and Pdx1 that changed the HPD cell fate to the intestinal epithelial cell fate that finally leads to form the intrahepatic intestinal tube in *hhex^zju1^* (Fig. 9). Pdx1 is a well-known factor for specifying the pancreas as well. The domain with ectopic expression of *pdx1* in *hhex^zju1^* did not become a part of pancreas, which suggests that Pdx1 alone is not sufficient to specify a pancreatic fate or Cdx1b or an unknown factor suppresses the Pdx1’s pancreatic function. Further investigation is needed to unravel the molecular mechanism behind.

**Fig. 9.**
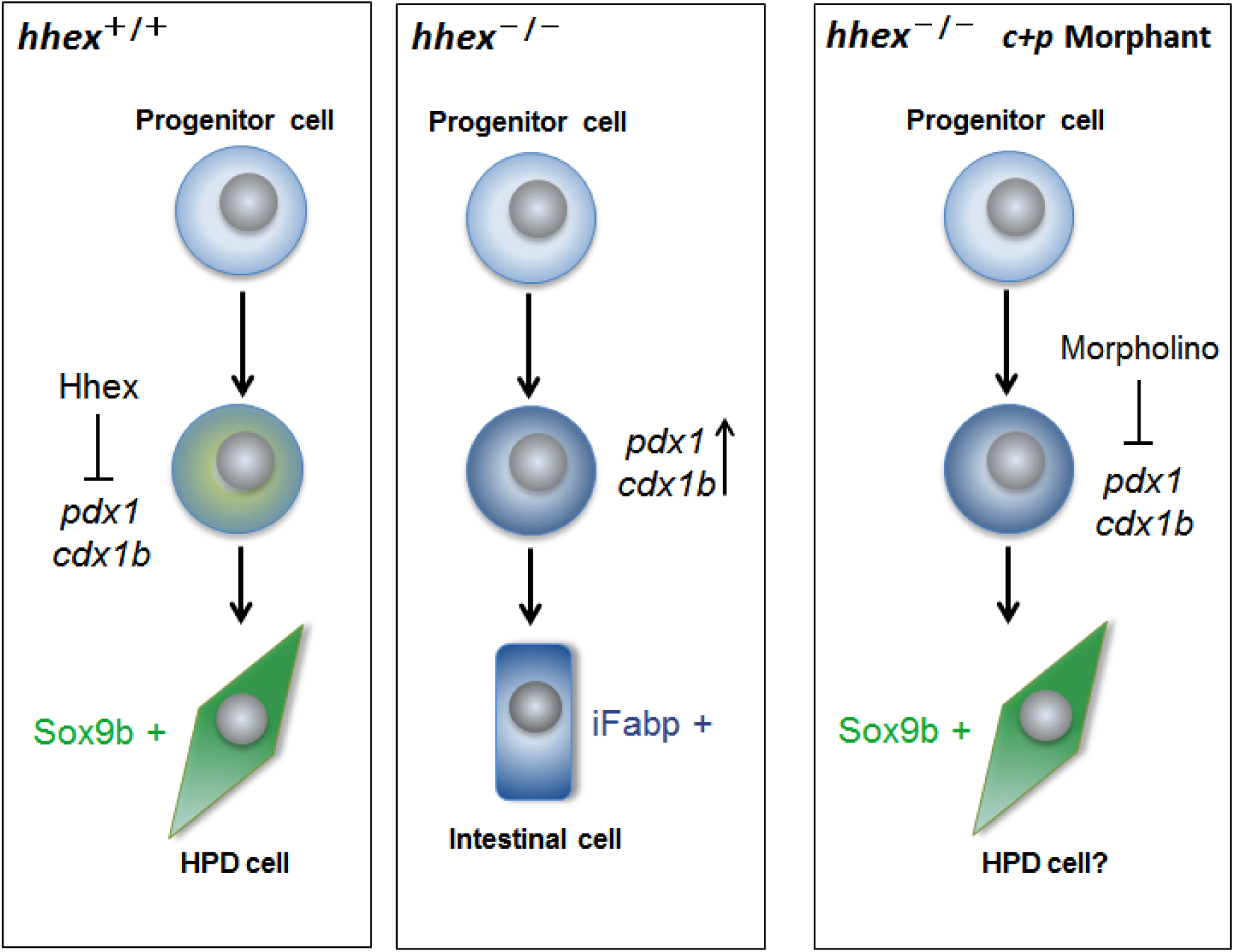
A model to depict the genetic network controlling the patterning and organogenesis of the HPD system. Based on the expression patterns of *hhex*, *cdx1b* and *pdx1* in WT and *hhex^zju1^* together with the analysis of the effect of Hhex on the promoters of *cdx1b* and *pdx1* genes, we propose that, in the WT HPD precursor cells, Hhex suppresses the expression of *cdx1b* and *pdx1* so that to safeguard the HPD cell fate. In *hhex^zju1^*, due to the absence of Hhex, the HPD precursor cells express Cdx1b and Pdx1 that changes the HPD cell fate to the intestinal epithelial cell fate that finally leads to the formation of an intrahepatic intestinal tube in *hhex^zju1^*. Morpholino-mediated knockdown of *cdx1b* and *pdx1* restores the *sox9b* expression in the HPD in *hhex^zju1^*. It appears that Cdx1b or Pdx1 or together, directly or indirectly suppresses the *sox9b* expression.

## MATERIALS AND METHODS

### Ethics statement

All animal procedures were performed in full accordance to the requirement by ‘Regulation for the Use of Experimental Animals in Zhejiang Province’. This work is approved by the Animal Ethics Committee in the School of Medicine, Zhejiang University (ETHICS CODE Permit NO. ZJU2011-1-11-009Y).

### Fish lines and maintenance

Zebrafish AB strain was used in all experiments and for generating transgenic or mutant lines. To generate zebrafish transgenic line *Tg(bhmt:EGFP)*, a 5.2kb genomic DNA fragment upstream of the start codon ATG of the *bhmt* gene is cloned with primer pair 5’-GCCATCCATGGCCCACATTCGT-3’ (forward primer) and 5’-GTTGATCTGATTCAGGAACAGCAGAT-3’ (reverse primer). The *bhmt:EGFP* construct was generated by subcloning EGFP cassettes downstream of the promoter sequences of *bhmt*. The *bhmt:EGFP* construct was linearized by *HindIII*, followed by injection into zebrafish embryos at the one-cell stage. The *Tg(bhmt:EGFP)* line was identified in the F1 generation. To generate *hhex* mutant, we synthesized gRNA against the first exon of zebrafish *hhex* gene according to the protocol described previously.(Chang et al. 2013) The *Cas9* mRNA and *hhex*-targeting gRNA was co-injected into the WT embryos at the one-cell stage. The *hhex* mutant lines were identified in the F1 generation by analyzing the PCR product using primer pair hhex ID (Supporting Table S1). For Cre/loxP-mediated lineage tracing, *Tg(fabp10a: CreERT2)* or *Tg(Tp1: CreERT2)*(He et al. 2014) embryos in the background of *Tg(β-actin:loxP-DsRed-STOP-loxP-GFP)* were treated with 10mM 4-hydroxytamoxifen (4-OHT) at 36 hpf for 3.5 days.

### WISH

The embryos were fixed in 4% PFA (PBS) for 12 h at 4°C. WISH Probes were labeled with digoxigenin (DIG, Roche Diagnostics). Probes *prox1*, *fabp10a*, *trypsin*, *insulin*, *fabp2a*, *foxa2*, *gata6* and *hhex* were used as described (Huang et al. 2008). For *cdx1b*, *sox9b* and *pdx1* probes, primers were designed based on available sequence data and PCR products were cloned into the pGEM-T Easy Vector (Promega) (Supporting Table S1).

### Cryosectioning and immunofluorescence staining

The embryos were fixed in 4% PFA (PBS) for 1 h at room temperature. After washing in PBST (0.1% Tween 20 in PBS), the tails of embryos were clipped for genomic DNA extraction for genotyping, and the rest parts were mounted with 1.5% agarose dissolved in 30% sucrose PBS and then equilibrated overnight at 4°C. The blocks were mounted with OCT compound (Sakura). The sections were cut serially to an 11-µm thickness and collected on poly-L-lysine coated glass slides (CITOGLAS, 188105). Immunofluorescence staining was performed as described.(Guan et al. 2016) Rabbit polyclonal antibody against zebrafish Fabp10a (1:500) and mouse monoclonal antibody against zebrafish Bhmt (1:500) were generated by Hangzhou HuaAn Biotechnology Company. P-H3 antibody was purchased from Santa Cruz (sc-8656-R, 1:600), PCNA antibody from Sigma (P8825, 1:1000) and 2F11 monoclonal antibody from Abcam (ab71286, 1:500). Alexa Fluor 647-labeled secondary antibody was used for visualization.

### RNA and protein analysis

RNA gel blot hybridization (northern blot) was performed as described (Huang et al. 2008). *EGFP* full length probe was DIG-labeled. Western blot was performed as described(Guan et al. 2016) using a mouse monoclonal antibody against zebrafish Hhex (1:200, HuaAn Biotechnology Company), anti-β-Actin antibody (1:1000, HuaAn R1207-1) and anti-EGFP antibody (1:1000, Santa Cruz sc-9996).

### Morpholino injection

*cdx1b* (1pmol) and *pdx1* (1pmol) ATG morpholino were co-injected into one-cell stage embryos. Morpholinos sequences were designed to target *cdx1b* ATG start codon site (5′-TCTAGGAGATAACTCACGTACATTT-3′) (Flores et al. 2008) or *pdx1* ATG codon start site (5’-GATAGTAATGCTCTTCCCGATTCAT-3’) (Kimmel et al. 2011).

### Luciferase assay

A series *cdx1b* or *pdx1* promoters were cloned in to the pGL3 vector with specific primer pairs (Table S1). Dual-luciferase reporter assays were carried out in cultured HepG2 cells or HCT116 cells. Cells were seeded in 24-well tissue culture plates 24 h prior to transfection. All transfections were performed in triplicate. For each well, co-transfection was carried out using 100ng of promoter assay plasmid (Firefly luciferase plasmids derived from pGL3), 30ng of either an empty expression vector (pCS2) or an expression vector encoding Hhex (pCS2-Hhex), 10ng of a control plasmid (Renilla luciferase vectors, phRL-TK) for normalizing the transfection efficiency, and 1μl PolyJet DNA Transfection Reagent (Signagen, SL100688) at 70–80% confluence. The cells were harvested 24h after transfection and assayed using the Dual-Luciferase^®^ Reporter Assay System (Promega, Cat.E1910).

### CHIP

ChIP was performed with slight modifications of the procedure described previously (Gong et al. 2015). HCT116 or HepG2 cells were seeded in 6cm tissue culture dishes 24 h prior to transfection. For each well, co-transfection was carried out using 500ng of promoter assay plasmid (Firefly luciferase plasmids derived from pGL3), 200ng of either a control expression vector encoding EGFP (pCS2-EGFP) or an expression vector encoding HA-Hhex (pCS2-HA-Hhex), 5μl PolyJet DNA Transfection Reagent (Signagen) at 70–80% confluence. At 24h post transfection, formaldehyde was added directly to tissue culture medium to a final concentration of 1% for 10 min at 37°C and was then stopped by the addition of glycine to a final concentration of 0.125 M. Cells were harvested and sonicated to shear DNA to lengths of about 300 base pairs. After centrifuging the samples, the supernatant was incubated overnight at 4°C with 20μl Anti HA-tag agarose beads (Abmart, M20013L). After serious extensively washing as described, the beads carried with antibody-protein–DNA complexes were added with proteinase-K reaction mix and heated at 65°C overnight to reverse protein–DNA crosslinks. The bound DNA fragments were purified by phenol/chloroform extraction and ethanol precipitation, and analyzed by Real-time Quantitative PCR (qPCR) Detecting System using specific primers (Table S2).

### Statistical analysis

Statistical analyses were performed with the Student’s T-test. * p < 0.05; ** p < 0.01; *** p < 0.001; n.s., no significant difference.

## Acknowledgments

The authors thank Drs Bin Zhao, Caiqiao Zhang, Hai Song and all members in JRP and CJ labs for their valuable suggestions. The authors are grateful to Yixin Ye, Xiaocai Du and Zhengxin Xu for their technical support with animal studies.

## SUPPLEMENTARY INFORMATION

**Figure S1.**
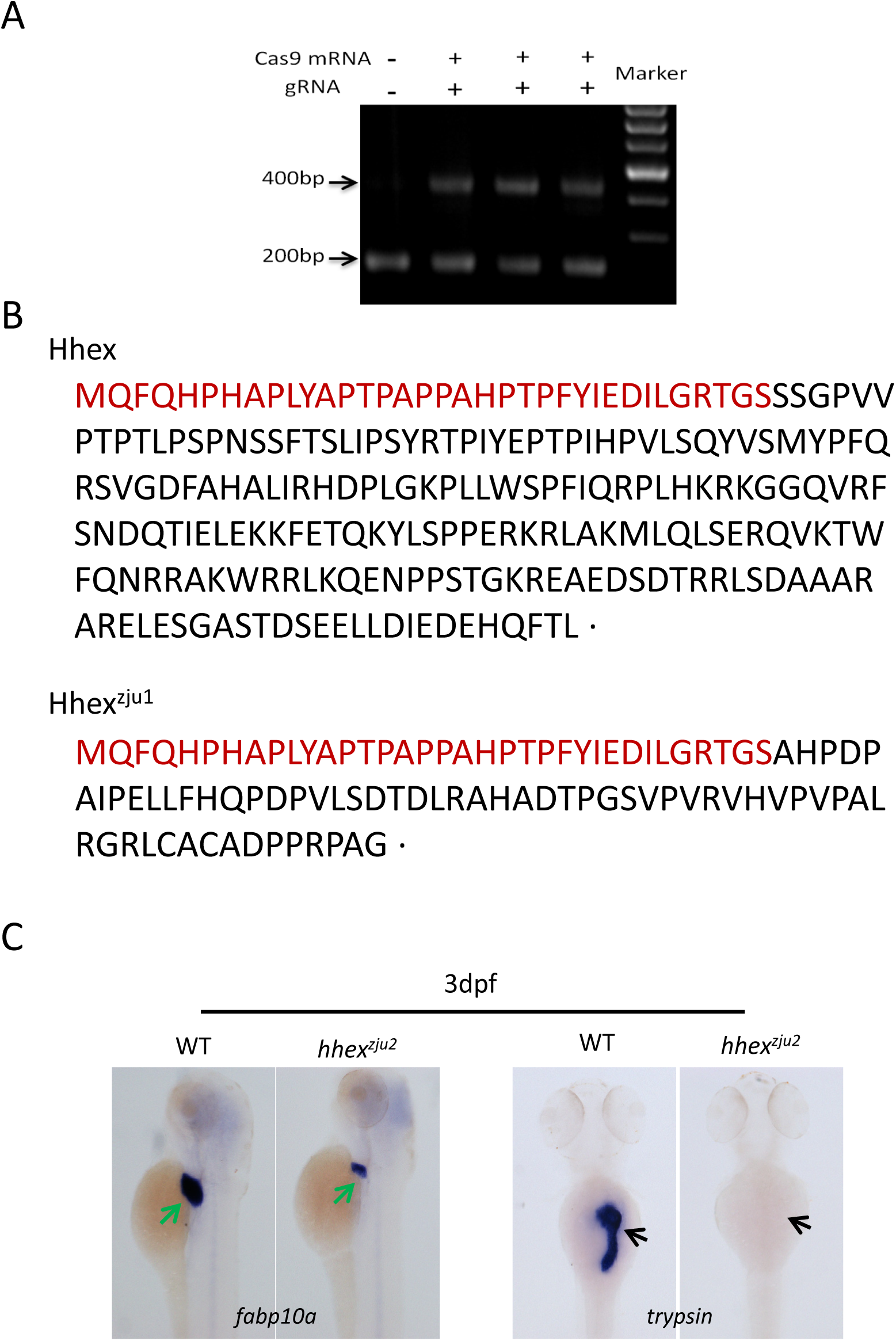
Generation of two *hhex* mutant alleles. (A) Gel photo showing the PCR products digested with *SacI* restriction enzyme. Three independent samples from 2.5dpf-old embryos co-injected with gRNA (+) and *cas9* (+) mRNA were shown. 400bp band: undigested product; 200bp band: digested product. (B) Upper panel: The amino acid (aa) sequence of the WT zebrafish Hhex protein. Lower panel: The *hhex^zju1^* mutation causes a frame shift to the WT *hhex* transcript and is predicted to produce a mutant peptide which only contains the first 35aa of the WT Hhex (in red letters). (C). WISH using the *fabp10a* (liver marker) or *trypsin* (exocrine pancreas marker) probes to show that the *hhex^zju2^* mutation also confers a small liver and near-absence of the pancreas phenotype as does the *hhex^zju1^* mutation.

**Figure S2.**
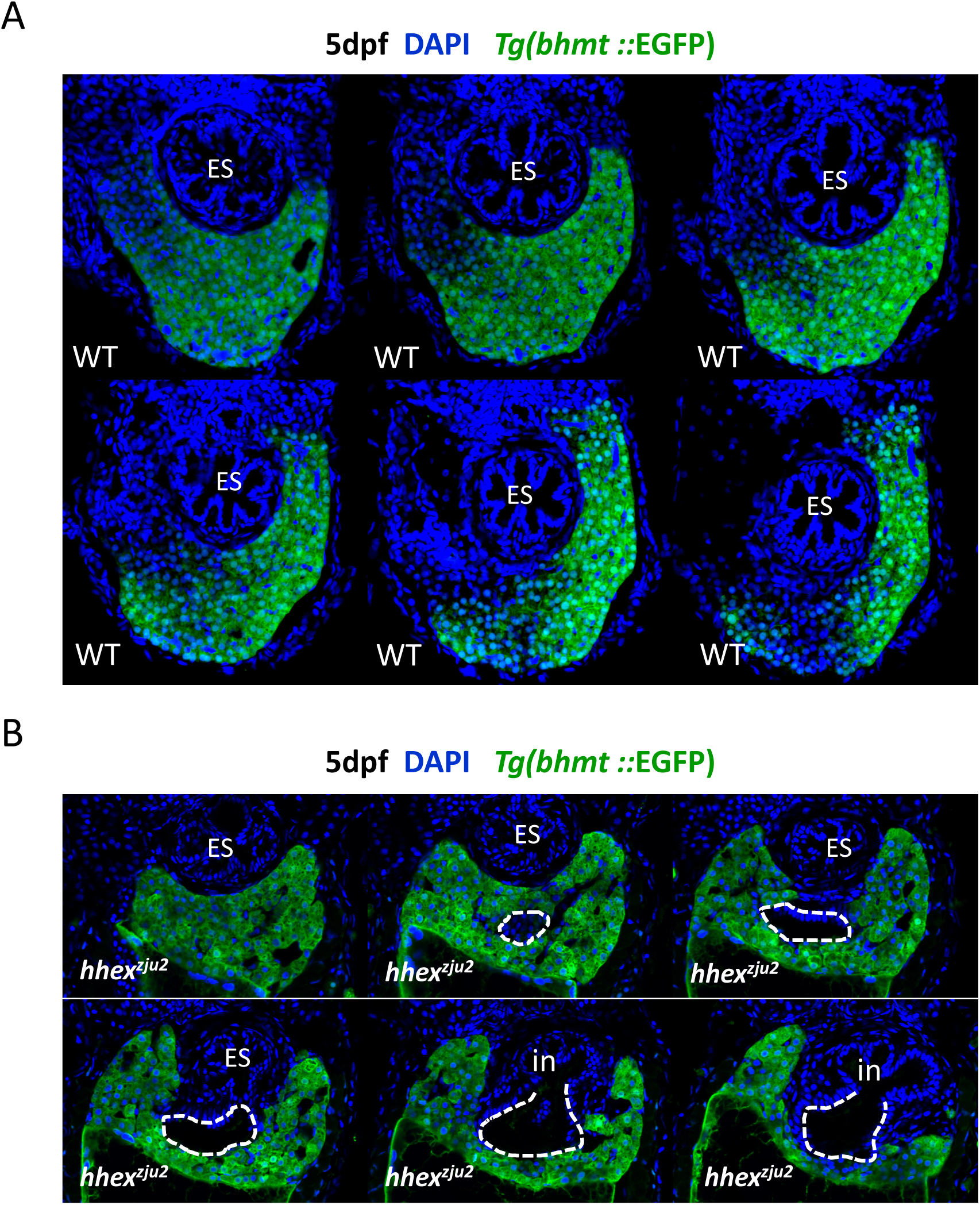
Loss-of-function of *hhex* leads to formation of an intrahepatic lumen structure. (A) Corresponding to Figure 3*A*, serial sections of a 5dpf-old WT embryo in the *Tg(bhmt:EGFP)* background show that there is no observable intrahepatic lumenized structure in the WT liver. (B) The *hhex^zju2^* mutant embryo (5dpf) develops an intrahepatic lumen (circulated with a white dashed line) in the liver that finally opens to the junction between the esophagus and intestinal tube as does the *hhex^zju1^* mutant shown in Figure 3*B*. ES: esophagus; in: intestine.

**Figure S3.**
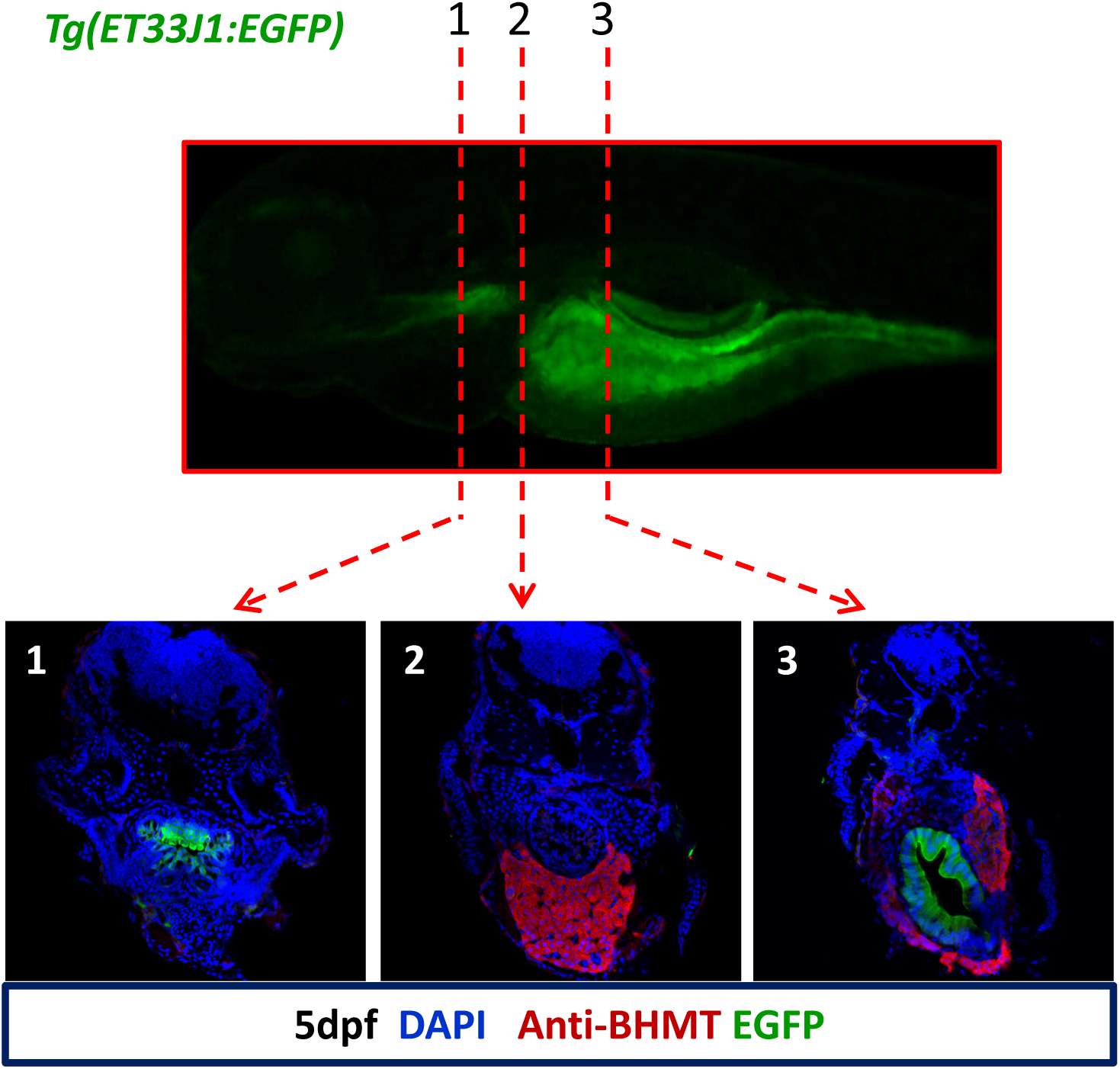
The *Tg(ET33J1:EGFP)* transgenic line is an intestinal epithelia reporter. Upper panel: an image showing the overall expression pattern of EGFP (green). Lower panels: Cross sections from the planes 1, 2 and 3 show that EGFP signals (green) can be observed in the anterior part of the esophagus (section 1) and the intestine tube (section 3) but not in the part of the esophagus (section 2) leading to the intestinal bulb. The Bhmt antibody was used to stain the hepatocytes (red) and DAPI was used to stain the nuclei (blue).

**Figure S4.**
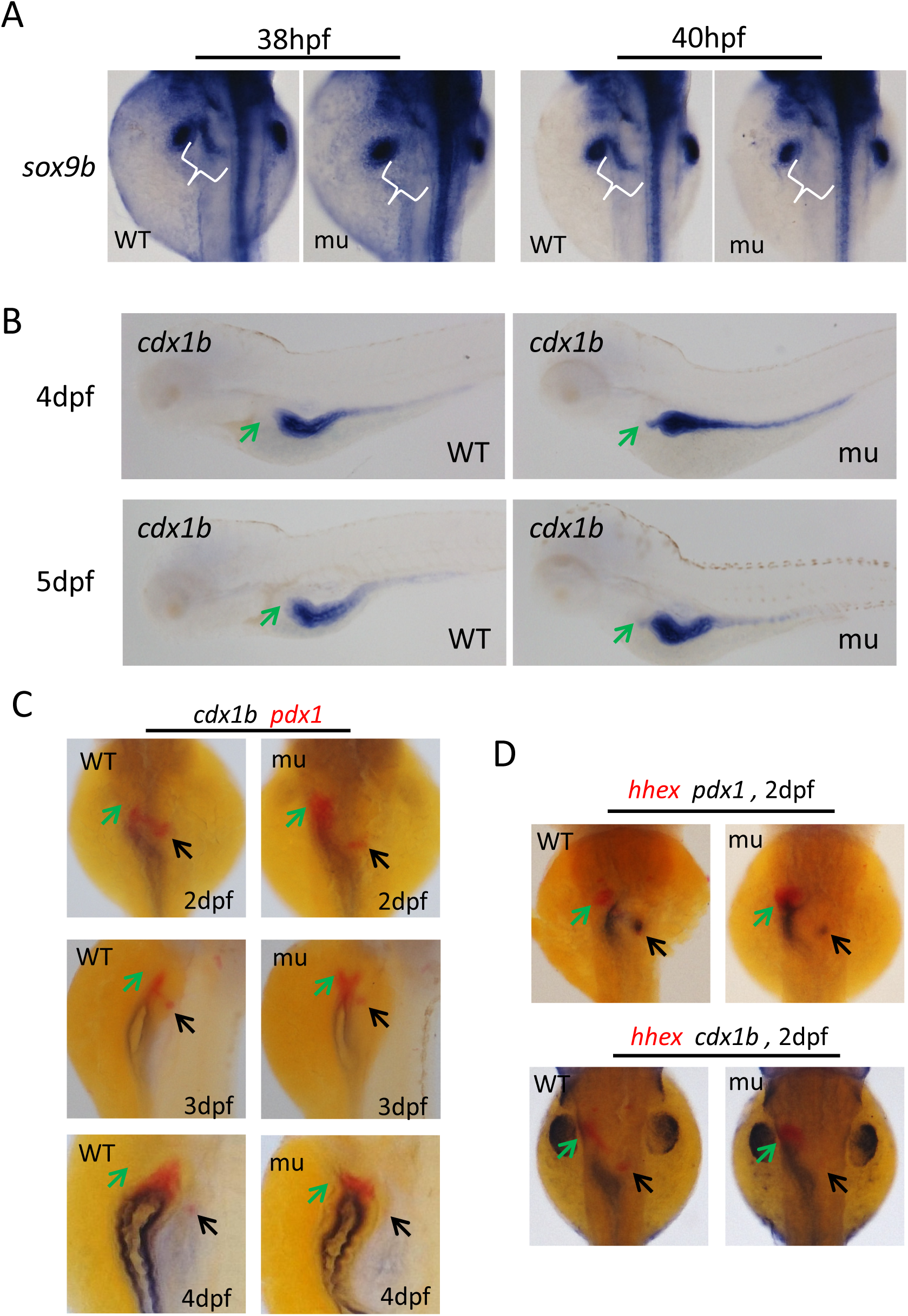
Loss-of-function of Hhex abolishes the *sox9b* expression in the HPD precursor cells but causes ectopic expression of *cdx1b* and *pdx1* in the liver bud. (A) WISH using the *sox9b* probe. In WT embryos (WT), *sox9b* transcripts are detected in the prospective HPD region (white bracket), head and the pectoral fin buds at 38hpf and 40hpf. In *hhex^zju1^* mutant embryos (mu), *sox9b* transcripts are also detected in the head and the pectoral fin buds, but not in the prospective HPD region (white bracket). (B) WISH using the *cdx1b* probe shows a protrusion of *cdx1b*-positive domain (yellow arrow) detoured to the liver bud from the anterior region of the foregut in the *hhex^zju1^* mutant at 4dpf and 5dpf, but that is never observed in a WT embryo. (C,D) WISH using *cdx1b* and *pdx1* double probes (C), *hhex* and *pdx1* (D, upper panels) or *hhex* and *cdx1b* (D, lower panels) double probes on WT and *hhex^zju1^* mutant (mu) embryos at 2dpf, 3dpf or 4dpf as shown. The middle line and somite 1 to 4 were indicated with yellow lines. Bracket, prospective HBPD; green arrow, liver; black arrow, islet; red arrow, the anterior edge of the intestinal bulb.

**Figure S5.**
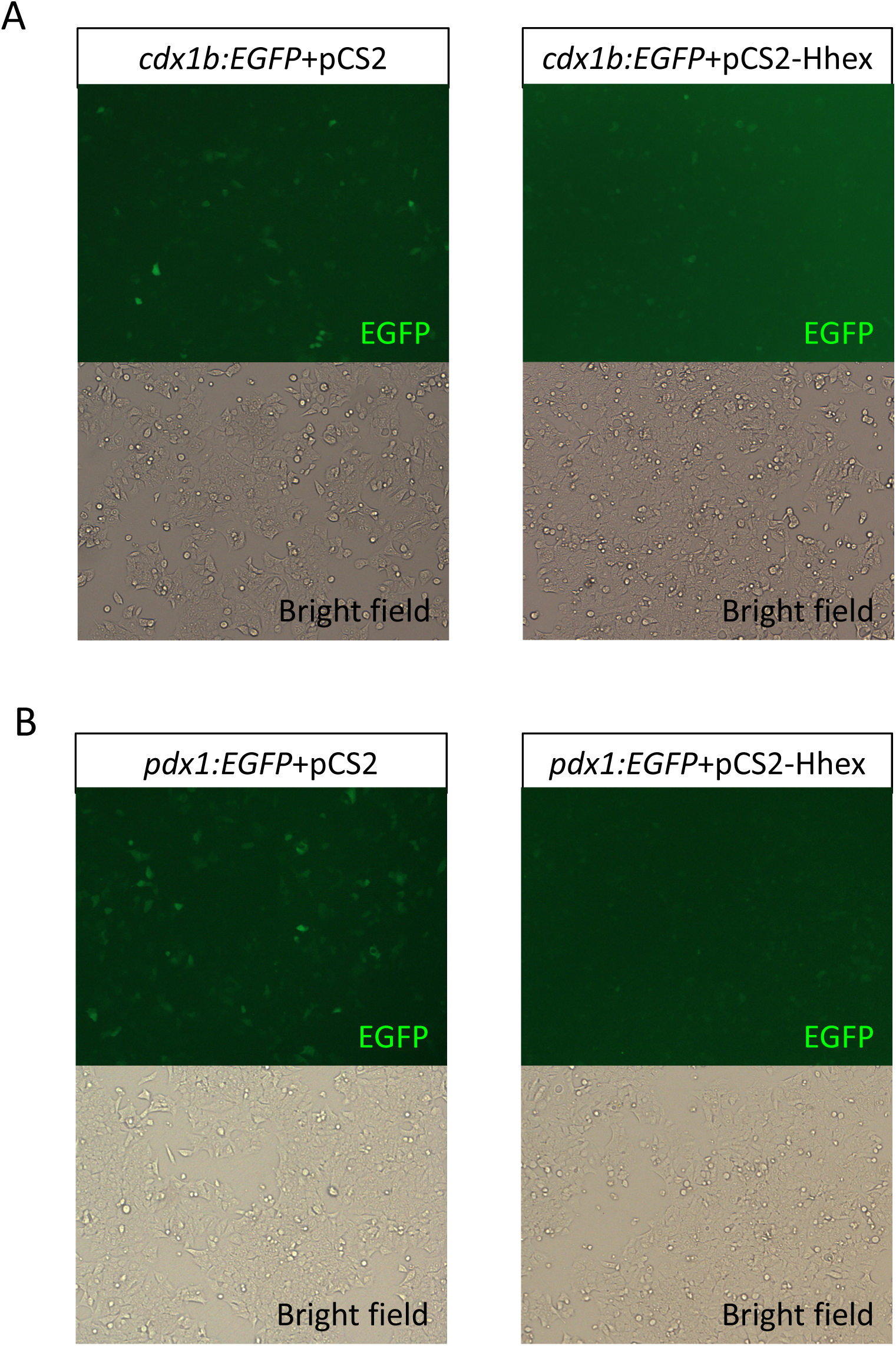
Hhex represses the transcriptional activity of *cdx1b* and *pdx1* gene promoters. (A,B). The *cdx1b:EGFP* (A) or *pdx1:EGFP* (B) plasmids was co-transfected with an empty expression vector (pCS2) or Hhex-expressing vector (*pCS2-Hhex*) into HCT116 cells. The EGFP fluorescence signal expressed by *cdx1b:EGFP* (A) or *pdx1:EGFP* (B) was greatly decreased in the cells expressing Hhex.

**Figure S6.**
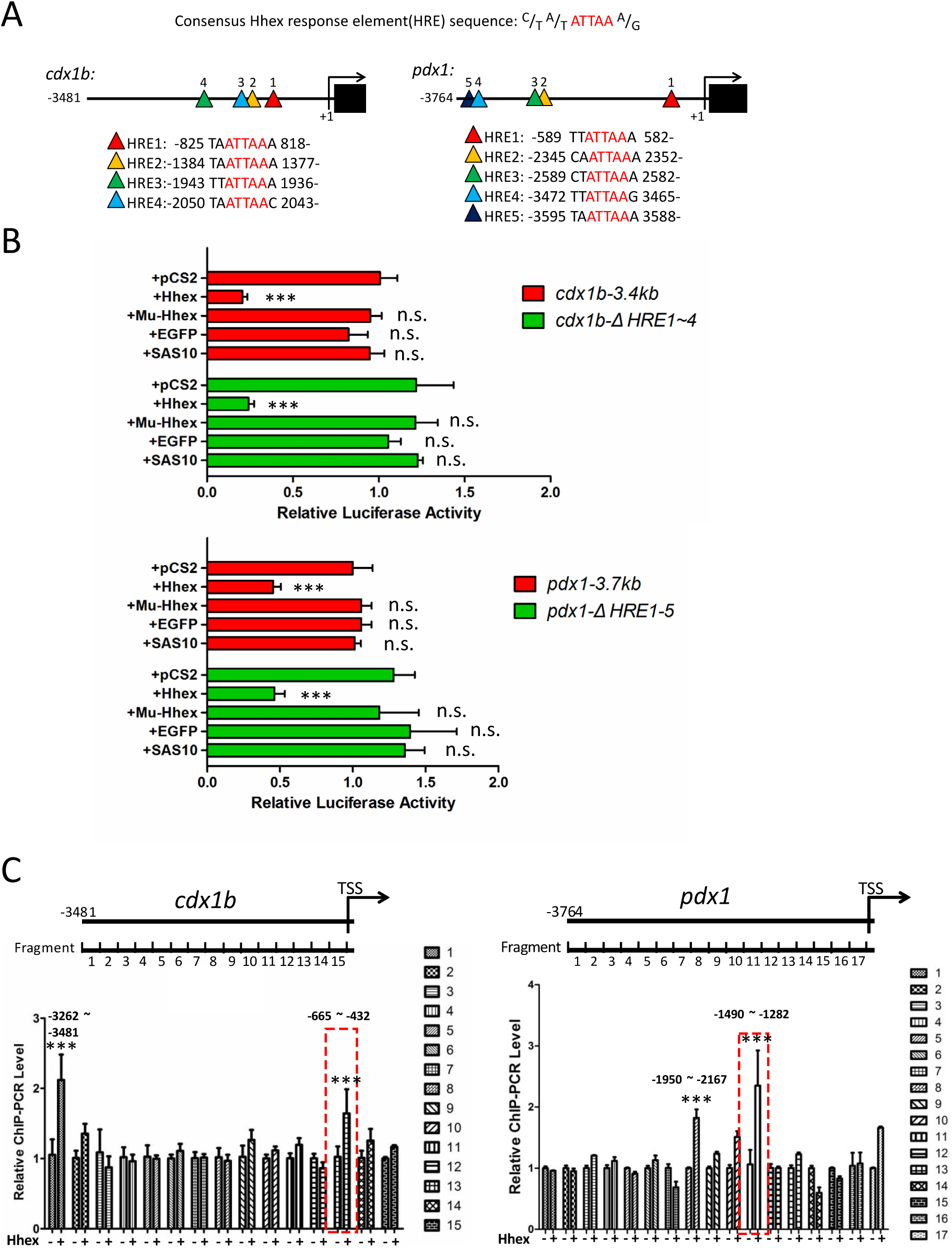
Identification of Hhex-response regions in *cdx1b* and *pdx1* promoters. (A). Upper panels: Diagrams showing the distribution and position of the putative Hhex-response element (HRE) in the promoters of *cdx1b* and *pdx1*. Each triangle represents one predicted HRE in the promoter fragment. Lower panels: Showing each of the putative HREs sequences and the five core nucleotides are highlighted in red. (B). Luciferase activity assay showed that deleting all four HREs in *cdx1b-3.4kb* (*cdx1b-ΔHRE1-4*) or all five HREs in *pdx1-3.7kb* (*pdx1-ΔHRE1-5*) exhibited similar responsiveness to Hhex as did the *cdx1b-3.4kb* and *pdx1-3.7* promoters. (C). The *cdx1b-3.4kb:luc* or *pdx1-3.7kb:luc* plasmid was co-transfected with or without Hhex-expressing plasmid (*pCS2-HA-Hhex*) into the HCT116 cells. Protein samples from such cells were subjected to ChIP assay using an antibody against the HA-tag to pulldown the Hhex-DNA hybrids. Fifteen (for *cdx1b-3.4kb*) and 17 (for *pdx1-3.7kb*) pairs of specific primers were designed, respectively, to amplify 15 and 17 DNA fragments covering the entire promoter sequences of *cdx1b* 3.481kb (left panels) and *pdx1* 3.764kb (right panels), respectively. The qPCR analysis of the ChIP product revealed that two segments in *cdx1b* (segments 1 and 13) and two fragments in *pdx1* (fragments 8 and 11) were enriched by HA-Hhex. * * *. P<0.001; n. s., no significance.

**Supporting Figure S7.**
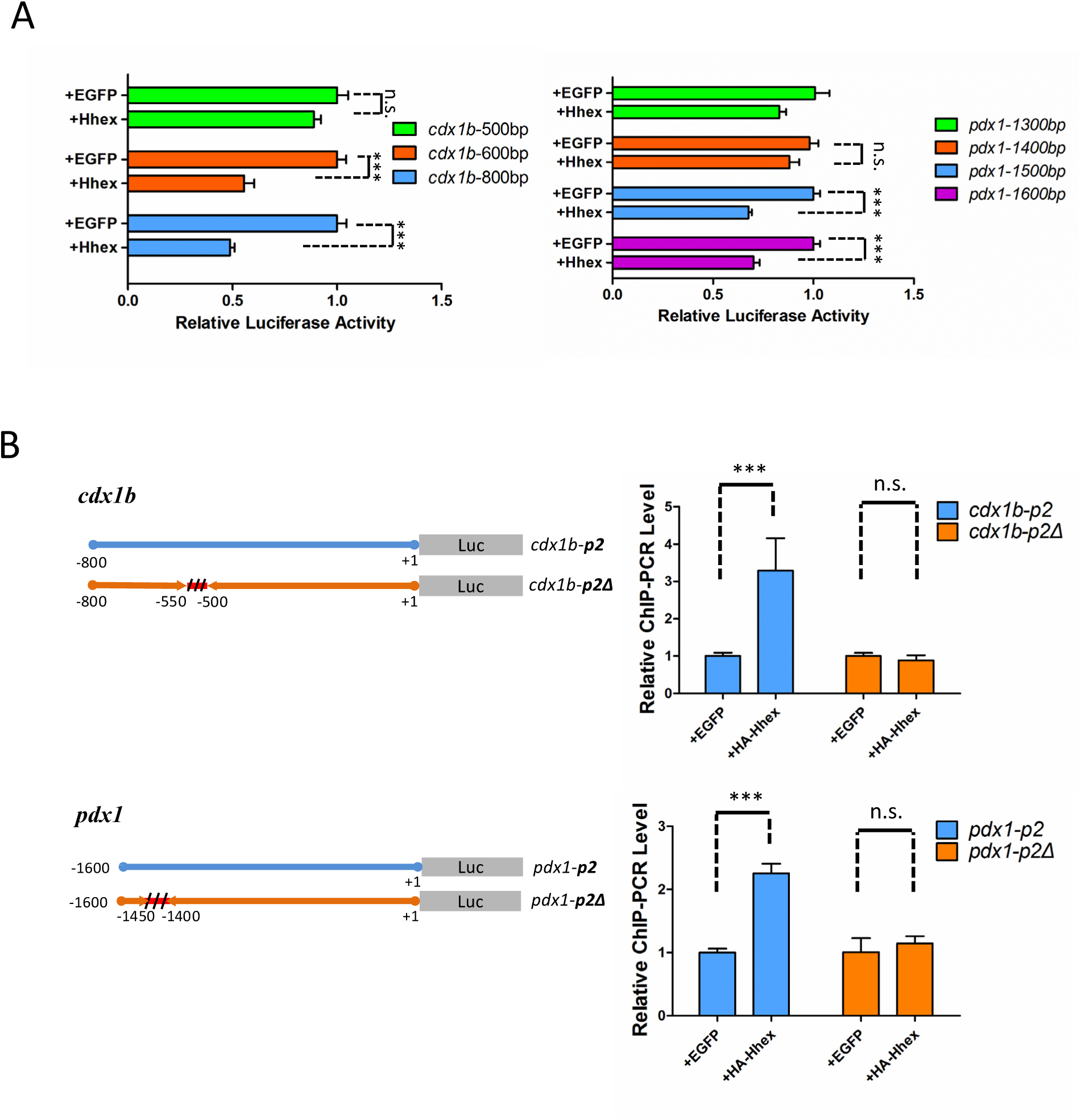
Identification of Hhex-response elements in *cdx1b* and *pdx1* promoters. (A). Based on the result shown in Supporting Figure S6C, the *cdx1b* promoter was truncated to generate three shortened promoter reporters, namely *cdx1b-*500bp (−1 to −500bp region), *cdx1b-*600bp (−1 to −600bp region) and *cdx1b*-800bp (−1 to −800bp region). Similarly, the *pdx1* promoter was truncated to generate four shortened promoter reporters, namely *pdx1-*1300bp (−1 to −1300bp region), *pdx1-*1400bp (−1 to −1400bp region), *pdx1*-1500bp (−1 to −1500bp region) and *pdx1-*1600bp (−1 to −1600bp region). Luciferase activity assay showed that *cdx1b-*600bp, *cdx1b*-800bp, *pdx1*-1500bp and *pdx1*-1600bp retained strong response to Hhex while *cdx1b-*500bp, *pdx1-*1300bp and *pdx1-*1400bp have lost the responsiveness to Hhex. (B). Corresponding to Figure 7*F* and *G*. The qPCR analysis of the ChIP products revealed that the internal deletion of −500 to −550bp in *cdx1b-p2* (*cdx1b*-800bp) (upper panels) and of −1400 to −1450bp in *pdx1-p2* (*pdx1*-1600bp) (lower panels) abolished the enrichment of *cdx1b-p2Δ* and *pdx1-p2Δ* by Hhex. ***. P<0.001; n.s., no significance.

**Supporting Table S1.**
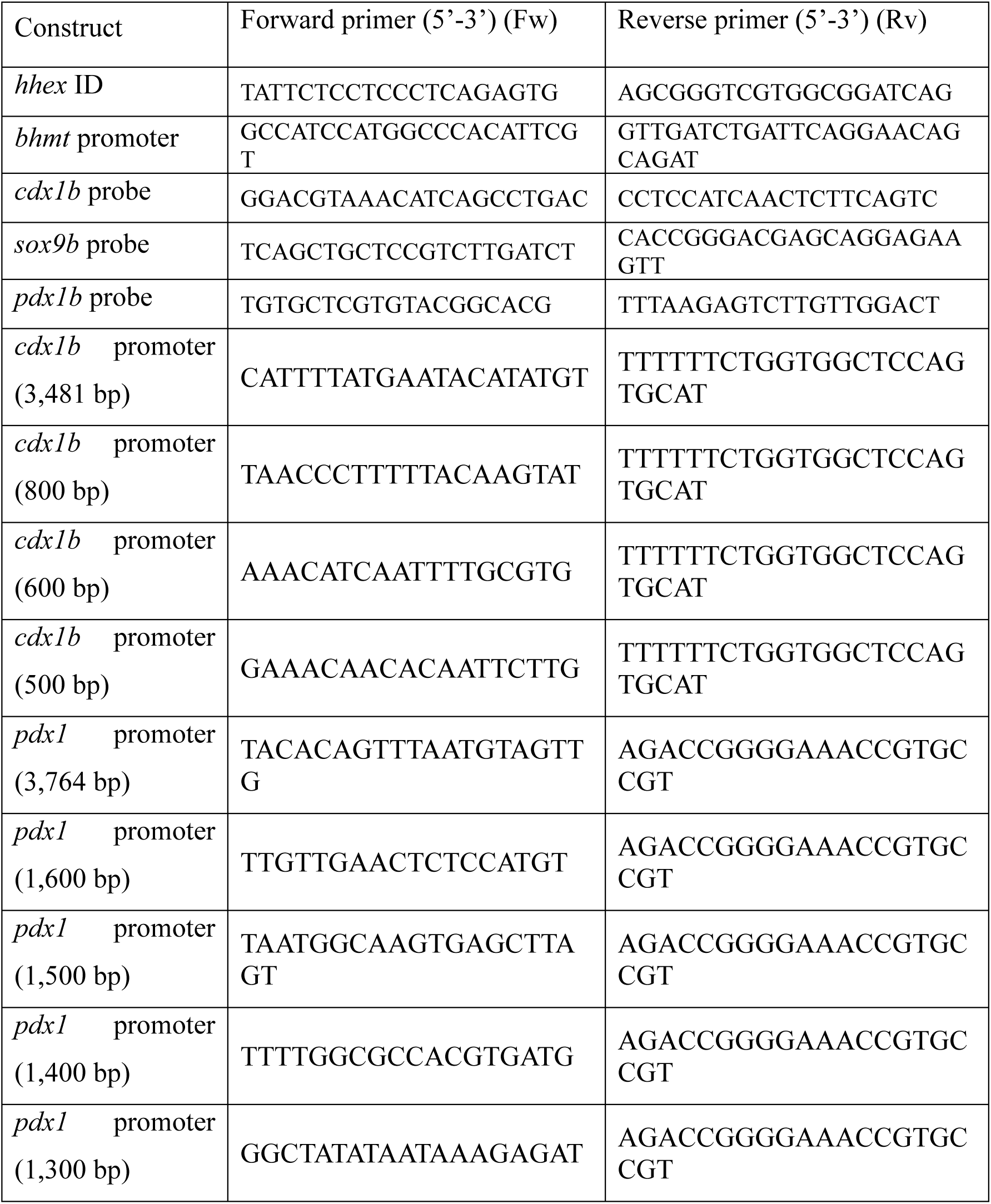
PCR primers used for cloning

**Supporting Table S2.**
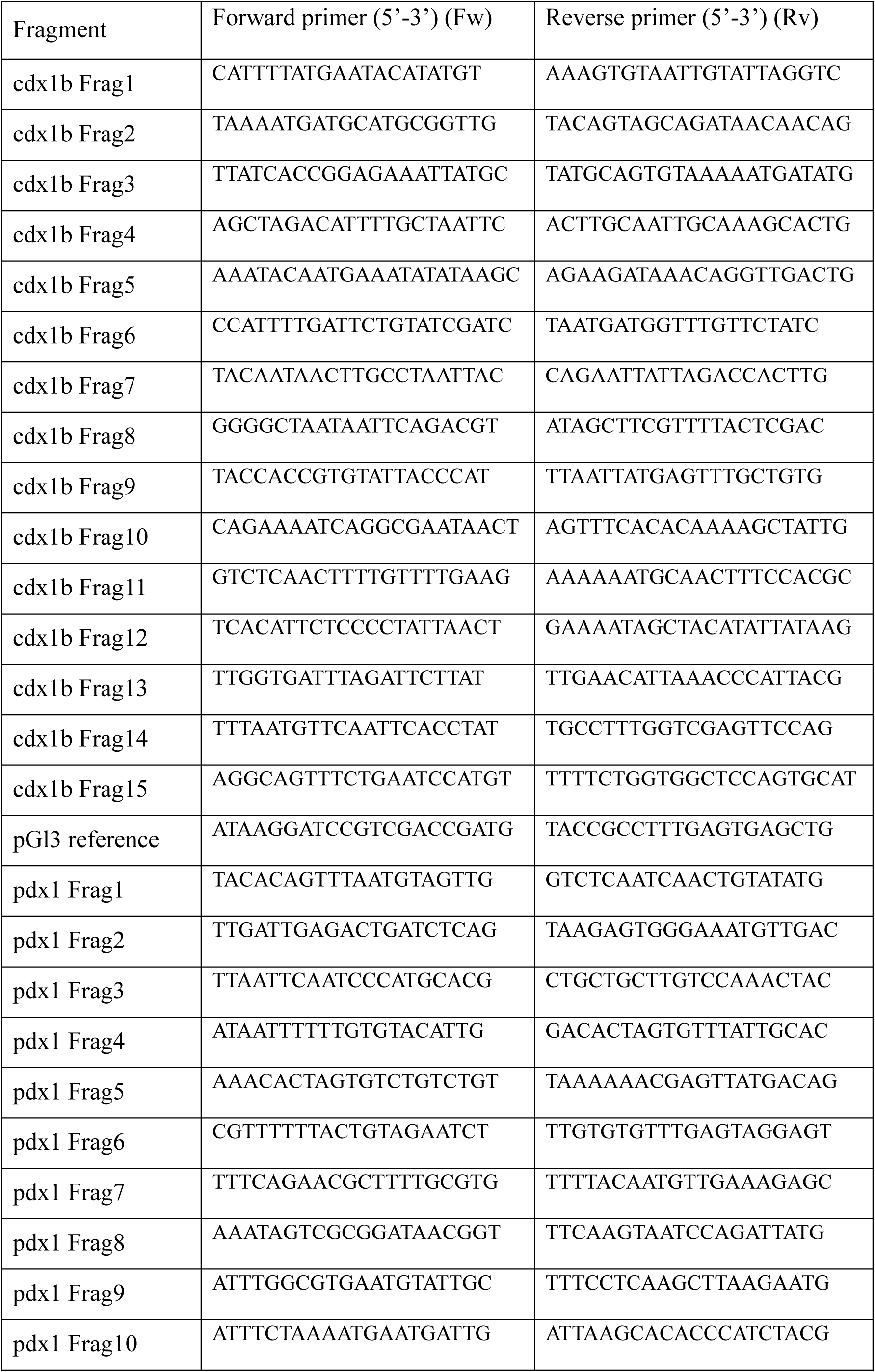
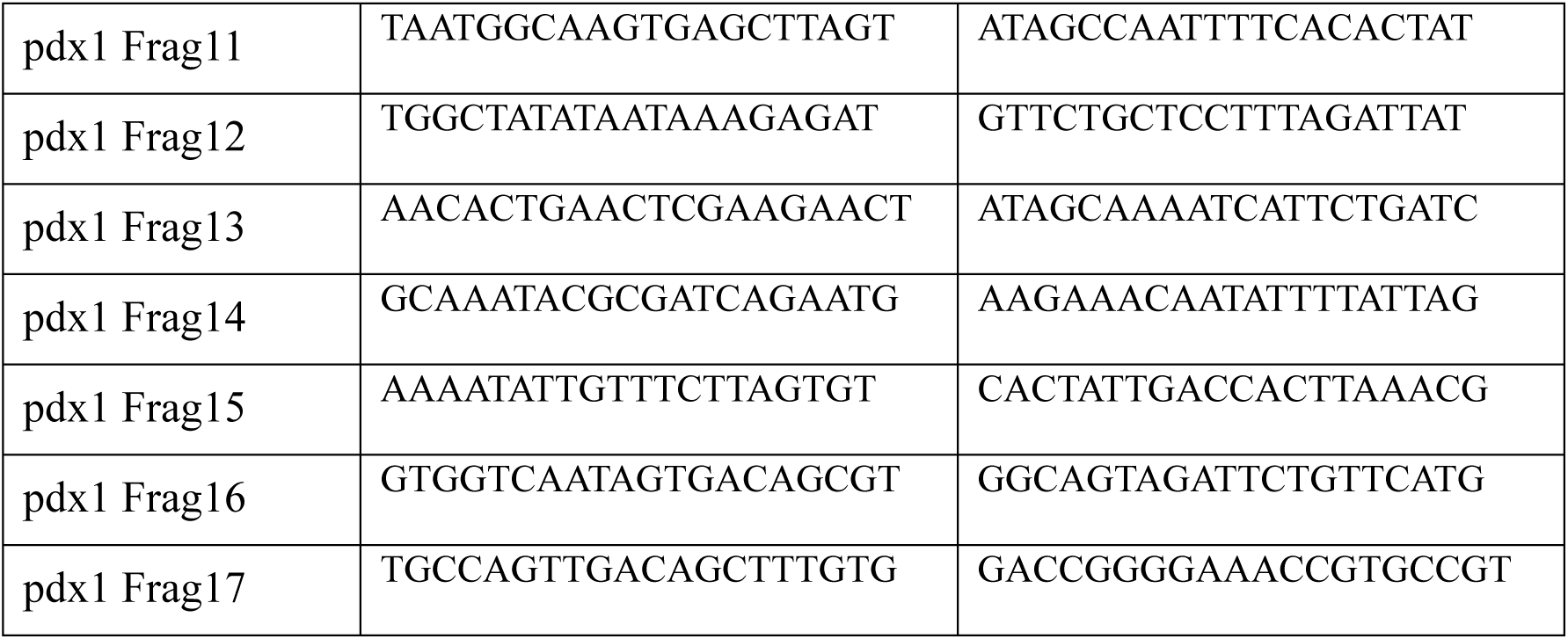
PCR primers used for ChIP PCR

## Abbreviations

Cdx1b: caudal-related homeobox 1b
ChIP: Chromatin-immunoprecipitation
HBPD: hepatobiliary and hepatopancreatic duct
Hhex: haematopoietically expressed homeobox
Pdx1: pancreatic and duodenal homeobox 1
WISH: whole-mount in situ hybridization

